# The bacterial chromatin protein HupA can remodel DNA and associates with the nucleoid in *Clostridium difficile*

**DOI:** 10.1101/426809

**Authors:** Ana M. Oliveira Paiva, Annemieke H. Friggen, Liang Qin, Roxanne Douwes, Remus T. Dame, Wiep Klaas Smits

## Abstract

The maintenance and organization of the chromosome plays an important role in the development and survival of bacteria. Bacterial chromatin proteins are architectural proteins that bind DNA, modulate its conformation and by doing so affect a variety of cellular processes. No bacterial chromatin proteins of *C. difficile* have been characterized to date.

Here, we investigate aspects of the *C. difficile* HupA protein, a homologue of the histone-like HU proteins of *Escherichia coli*. HupA is a 10 kDa protein that is present as a homodimer *in vitro* and self-interacts *in vivo*. HupA co-localizes with the nucleoid of *C. difficile*. It binds to the DNA without a preference for the DNA G+C content. Upon DNA binding, HupA induces a conformational change in the substrate DNA *in vitro* and leads to compaction of the chromosome *in vivo*.

The present study is the first to characterize a bacterial chromatin protein in *C. difficile* and opens the way to study the role of chromosomal organization in DNA metabolism and on other cellular processes in this organism.

## Introduction

*Clostridium difficile* (also known as *Clostridioides difficile*)[1] is a gram-positive anaerobic bacterium that can be found in environments like soil, water, and even meat products [2, 3]. It is an opportunistic pathogen and the leading cause of antibiotic-associated diarrhea in nosocomial infections [4]. *Clostridium difficile* infection (CDI) can present symptoms that range from mild diarrhea to more severe disease, such as pseudomembranous colitis, and can even result in death [4]. Over the past two decades the incidence of CDI worldwide, in a healthcare setting as well as in the community has increased [4–6]. *C. difficile* is resistant to a broad range of antibiotics and recent studies have reported cases of decreased susceptibility of *C. difficile* to some of the available antimicrobial therapies [7, 8]. Consequently, the interest in the physiology of the bacterium has increased in order to explore new potential targets for intervention.

The maintenance and organization of the chromosome plays an important role in the development and survival of bacteria. Several proteins involved in the maintenance and organization of the chromosome have been explored as potential drug targets [9–11]. The bacterial nucleoid is a highly dynamic structure organized by factors such as the DNA supercoiling induced by the action of topoisomerases [12], macromolecular crowding [13, 14] and interactions with nucleoid-associated proteins (NAPs) [15, 16]. Bacterial NAPs have been implicated in efficiently compacting the nucleoid while supporting the regulation of specific genes for the proliferation and maintenance of the cell [16].

NAPs are present across all bacteria and several major families have been identified [16, 17]. Some of the most abundant NAPs in the bacterial cell are bacterial chromatin proteins like the HU/IHF protein family [18, 19]. *Escherichia coli* contains three HU/IHF family proteins (αHU, βHU, IHF) that have been extensively characterized [19–22]. By contrast, *Bacillus subtilis* and several other gram-positive organisms only contain one protein of the HU/IHF protein family [17, 19, 23]. In *E. coli* disruption of αHU and/or βHU function leads to a variety of growth defects or sensitivity to adverse conditions, but HU is not essential for cell survival [24, 25]. However, in *B. subtilis* the HU protein HBsu is essential for cell viability, likely due to the lack of functional redundancy of the HU proteins such as in *E. coli* [17, 23].

In solution, most HU proteins are found as homodimers or heterodimers and are able to bind DNA through a flexible DNA binding domain. The crystal structure of the *E. coli* αHU-βHU heterodimer suggests the formation of higher order complexes at higher protein concentrations [22]. Modeling of these complexes suggest HU proteins have the ability to form higher-order complexes through dimer-dimer interaction and make nucleoprotein filaments [22, 26, 27]. However, the physiological relevance of these is still unclear [18, 22, 27].

The flexible nature of the DNA-binding domain in HU proteins confers the ability to accommodate diverse substrates. Most proteins bind with variable affinity and without strong sequence specificity to both DNA and RNA [28]. Some bacterial chromatin proteins have a clear preference for AT-rich regions [29–31] or for the presence of different structures on the DNA [28, 32].

HU proteins can modulate DNA topology in various ways. They can stabilize negatively supercoiled DNA or constrain negative supercoils in the presence of topoisomerase [22, 33]. HU proteins are involved in modulation of the chromosome conformation and have been shown to compact DNA [16, 26, 34]. This compaction of DNA is possible through the ability of HU proteins to introduce flexible hinges and/or bend the DNA [16, 26, 34, 35].

The ability to induce conformational changes in the DNA influences a variety of cellular processes due to an indirect effect on global gene expression [36–40]. In *E. coli* HU proteins are differentially expressed during the cell cycle. The αHU-βHU heterodimer is prevalent in stationary phase, while during exponential growth HU is predominantly present as homodimers [21]. Several studies suggest an active role of HU proteins in the transcription and translation of other proteins and even in DNA replication and segregation of the chromosomes [41–43].

The diverse roles of HU proteins are underscored by their importance for metabolism and virulence in bacterial pathogens. Disruption of both HU homologues (αHU and βHU) in *Salmonella typhimurium*, for example, results in the down-regulation of the pathogenicity island SPI2 and consequently a reduced ability to survive during macrophage invasion [44]. Other studies have shown the importance of HU proteins for the adaptation to stress conditions, such as low pH or antibiotic treatment [45–47]. For instance, in *M. smegmatis* deletion of *hupB* leads to increased sensitivity to antimicrobial compounds [46].

Despite the wealth of information from other organisms, no bacterial chromatin protein has been characterized to date in the gram-positive enteropathogen *Clostridium difficile*. In this study, we show that *C. difficile* HupA (CD3496) is a legitimate homologue of the bacterial HU proteins. We show that HupA exists as a homodimer, binds to DNA and co-localizes with the nucleoid. HupA binding induces a conformational change of the substrate DNA and leads to compaction of the chromosome. This study is the first to characterize a bacterial chromatin protein in *C. difficile*.

## Results and Discussion

### C. difficile *encodes a single HU protein, HupA*

To identify bacterial chromatin proteins in *C. difficile*, we searched the genome sequence of *C. difficile* for homologues of characterized HU proteins from other organisms. Using BLASTP (https://blast.ncbi.nlm.nih.gov/), we identified a single homologue of the HU proteins in the genome of the reference strain 630 [48]; GenBank: AM180355.1), encoded by the *hupA* gene (CD3496)(e-value: 1e-22). This is similar to other gram-positive organisms, where also a single member of this family is found [17, 19, 23] and implies an essential role of this protein on the genome organization in *C. difficile*. Moreover, lack of *hupA* mutants during random transposon mutagenesis of the epidemic *C. difficile* strain R20291 supports that the *hupA* gene (CDR20291_3333) is essential [49].

Alignment of the HupA amino acid sequence with selected homologues from other organisms reveals a sequence identity varying between 58% to 38% (Fig. 1a). HupA displays the highest sequence identity with *Staphylococcus aureus* HU (58%). When compared to the *E. coli* HU proteins, HupA has a higher sequence identity with βHU (47%) than with αHU (43%).

**Fig. 1.**
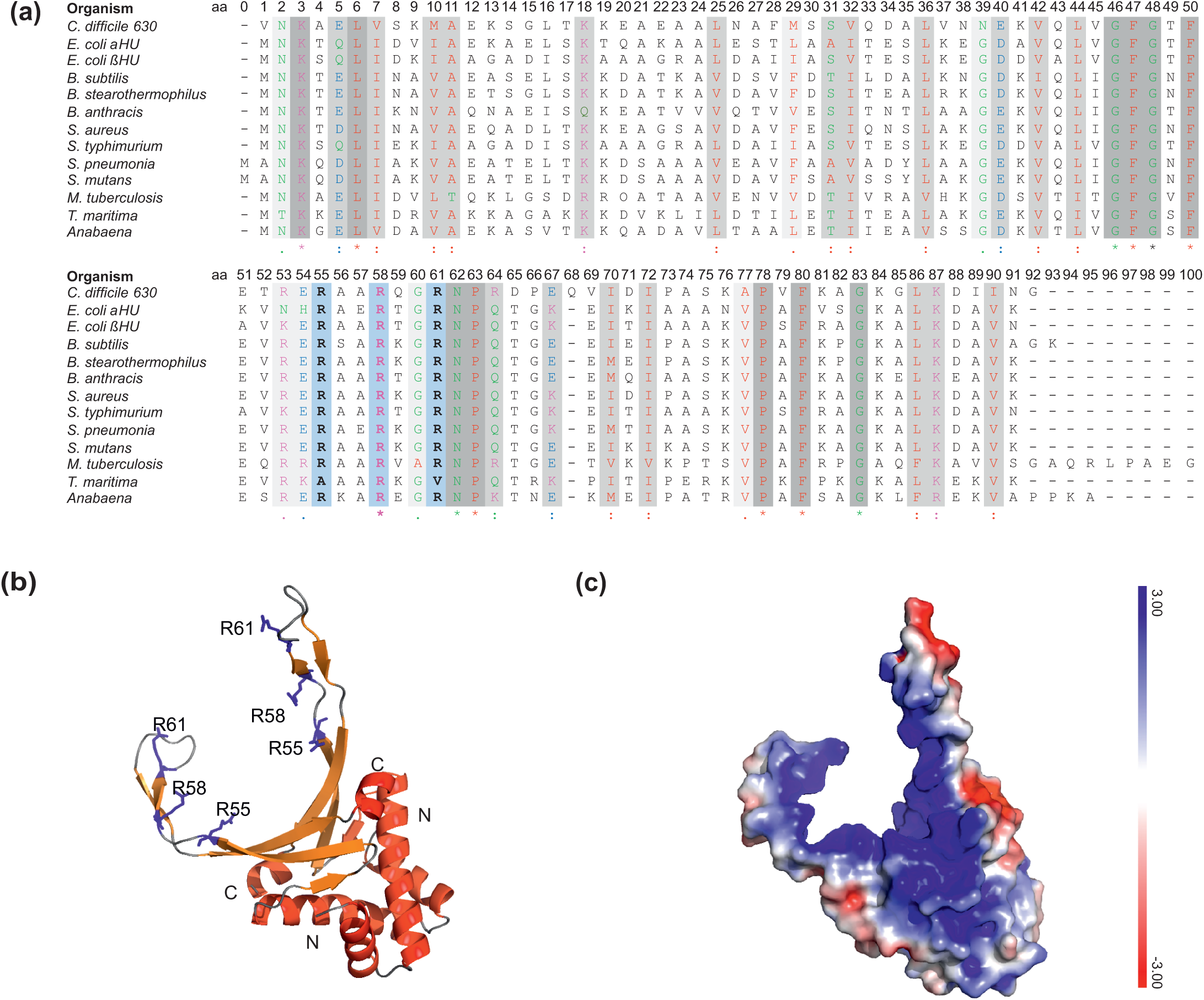
*Clostridium difficile* HupA is a homologue of bacterial HU proteins. (a) Multiple sequence alignment (ClustalOmega) of *C. difficile* HupA with homologous proteins from the Uniprot database. The protein sequences from *C. difficile* 630Δ*erm* (Q180Z4), *Escherichia coli* αHU (P0ACF0), *E. coli* βHU (P0ACF4), *Bacillus subtilis* (A3F3E2), *Geobacillus stearothermophilus* (P0A3H0), *Bacillus anthracis* (Q81WV7), *Staphylococcus aureus* (Q99U17), *Salmonella typhimurium* (P0A1R8), *Streptococcus pneumoniae* (AAK75224), *Streptococcus mutans* (Q9XB21), *Mycobacterium tuberculosis* (P9WMK7), *Thermotoga. maritima* (P36206) and *Anabaena sp.* (P05514) were selected for alignment. Residues are coloured according to ClustalW2 convention. Conserved residues (indicated with symbols below the alignment) are additionally highlighted with grey shading (darker = more conserved), except for the three arginine residues that were subjected to mutagenesis (in bold), that are highlighted in blue. (b) Structural model of the *C. difficile* HupA dimer based on homology with the crystal structure of DNA-bound nucleoid associated protein SAV1473 (SWISS-MODEL, PDB: 4QJN, 58.43% identity). α-helixes are represented in red, β-sheets in orange and unstructured regions in grey. Both the N-terminus and the C-terminus are indicated in the figure. A DNA binding pocket is formed by the arm regions of the dimer, composed by 4 β-sheets in each monomer. The localization of the substituted residues (R55, R58 and R61) are indicated (blue, sticks). (c) Electrostatic surface potential of *C. difficile* HupA. The electrostatic potential is in eV with the range shown in the corresponding color bar.

The overall structure of HU proteins is conserved and has previously been described through the analysis of several nucleoid-associated proteins [19, 50]. To confirm the structural similarity of the *C. difficile* HupA protein to other HU proteins, we performed a PHYRE2 structure prediction [51]. All top-scoring models are based on structures from the HU family. The model with the highest confidence (99.9) and largest % identity (60%) is based on a structure of the *S. aureus* HU protein (PDB: 4QJU). Next, we generated a structural model of HupA using SWISS-MODEL [52] and *S. aureus* HU protein (Uniprot ID: Q99U17)[53] as a template. As expected, the predicted structure (Fig. 1b) is a homodimer, in which each monomer contains two domains as is common for HU proteins [50, 53]. The α-helical dimerization domain contains a helix-turn-helix (HTH) and the DNA-binding domain consists of a protruding arm composed of 3 β-sheets (Fig. 1b). In the dimer, the two β-arms form a conserved pocket that can extensively interact with the DNA [53](Fig 1a).

Crystal structures of HU-DNA complexes have shed light on the mode of interaction of HU proteins with DNA [35, 53–55]. In the co-crystal structure of *S. aureus* HU the arms embrace the minor groove of the dsDNA [53]. Proline residues at the terminus of the arms cause distortion of the DNA helix, by creating or stabilizing kinks [35, 53]. Further electrostatic interactions between the sides of HU dimers and the phosphate backbone facilitate DNA bending [56]. In *Borrelia burgdorferi* direct interactions between the DNA backbone and the α-helices of the Hbb protein dimerization domain were observed [55]. The overall similarity of *C. difficile* HupA to other HU family proteins (Fig. 1a) and a similar predicted electrostatic surface potential (Fig. 1c) suggest a conserved mode of DNA binding for *C. difficile* HupA.

### Mutating arginine residues in the beta-arm of HupA eliminates DNA binding

Based on the alignment and structural model of HupA (Fig.1) we predict that several amino acid residues in *C. difficile* HupA could be involved in the interaction with DNA. Specifically, the positively charged arginine residues R55, R58 and R61 on the β-arms of HupA (Fig. 1a and b) were of interest. In *B. stearothermophilus* arginine 55 of BstHU (residue reference to *C. difficile*) is essential for the interaction with DNA, while residues R58 and R61 have a minor effect [57]. In contrast, R58 and R61 play an important role in DNA binding of *E. coli* βHU [58]. In *S. aureus* substitutions of the residue R58, reduced the affinity of HU for DNA while R55 and R61 were crucial for proper DNA binding [53].

As it has been shown that disruption of a single residue may not be sufficient to abolish DNA binding [32, 57, 58], we substituted the residues R55, R58 and R61 (Fig. 1b, blue sticks) in *C. difficile* HupA based on the published mutations in HU from other organisms [53, 57, 58]. Residue R55 was changed to glutamine (Q), a neutral residue with long side chain. R58 and R61 were replaced by glutamic acid (E) and aspartic acid (D), respectively, both negatively charged residues. The resulting protein is referred to as HupA^QED^. Evaluation of the effect of these mutations on the electrostatic surface potential of the structural model of HupA reveals that compared to the wild-type protein (Fig. 1c), HupA^QED^ exhibits a reduced positively charged surface of the DNA binding pocket (Fig. S1), which is expected to prevent the interaction with DNA.

To test the DNA binding of HupA and HupA^QED^ we performed gel mobility shift assays. *C. difficile* HupA and HupA^QED^ were heterologously produced and purified as 6x histidine-tagged fusion proteins (HupA_6xHis_ and HupA^QED^_6xHis;_ see Materials and Methods). We incubated increasing concentrations of protein with different [γ-^32^P]-labelled 38 bp double-stranded DNA (dsDNA) fragments with different G+C content. When HupA_6xHis_ was incubated with the DNA fragment a progressive reduction in mobility as a function of protein concentration is evident (Fig. 2a). At 2 µM of protein, approximately 70% of DNA is present as a DNA:protein complex (Fig. 2b). This clearly demonstrates that HupA_6xHis_ is capable of interacting with DNA.

**Fig. 2.**
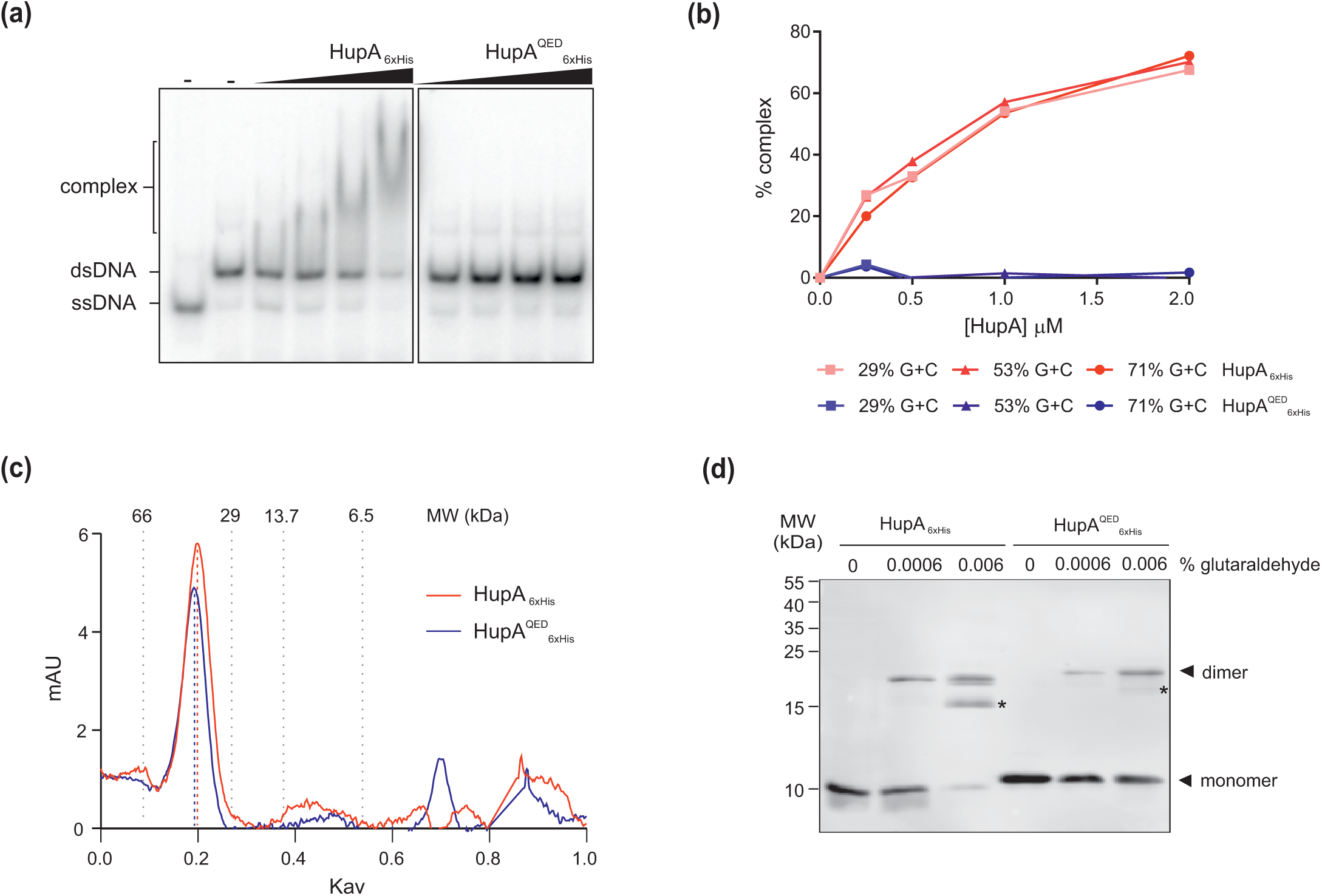
Dimerization of HupA is independent of DNA binding. (a) Electrophoretic mobility shift assays with increasing concentrations (0.25 −2 µM) of HupA_6xHis_ and HupA^QED^_6xHis_. Gel shift assays were performed with 2.4 nM radio-labeled ([γ-^32^P] ATP) 29% G+C dsDNA oligonucleotide incubated with HupA for 20 min at room temperature prior to separation. Protein-DNA complexes were analysed on native 8% polyacrylamide gels, vacuum-dried and visualized by phosphorimaging. ssDNA and dsDNA (without protein added, “-”) were used as controls. (b) Quantification of the gel-shift DNA-protein complex by densitometry. Gel shift assays were performed with 2.4 nM radio labelled ([γ-^32^P] ATP) dsDNA oligonucleotides with different 29-71% G+C content and the indicated concentration of HupA_6xHis_ (red) and HupA^QED^_6xHis_ (blue). (c) Elution profiles of HupA_6xHis_ (red) and HupA^QED^_6xHis_ (blue) from size exclusion chromatography. The experiments were performed with purified protein (100 μM) on a Superdex HR 75 10/30 column. The elution position of protein standards of the indicated MW (in kDa) are indicated by vertical grey dashed lines. The elution profiles show a single peak, corresponding to a ∼38 kDa multimer, when compared to the predicted molecular weight of the monomer (11 kDa). No significant difference in the elution profile of the HupA^QED^_6xHis_ compared to HupA_6xHis_ was observed. (d) Western-blot analysis of glutaraldehyde cross-linking of HupA_6xHis_ and HupA^QED^_6xHis_. 100 ng HupA was incubated with 0%, 0.0006% and 0.006% glutaraldehyde for 30 min at room temperature. The samples were resolved by SDS-PAGE and analysed by immunoblotted with anti-his antibody. Crosslinking between the HupA monomers is observed with the approximate molecular weight of a homo-dimer (∼22 kDa). Additional bands of lower molecular weight HupA are observed (*) that likely represent breakdown products.

Some nucleoid-associated proteins demonstrate a preference for AT-rich regions [29, 30, 59]. We considered that binding of HupA could show preference for low G+C content DNA, since *C. difficile* has a low genomic G+C content (29.1% G+C). We tested DNA binding to dsDNA with 71.1%; 52.6% and 28.9% G+C content but observed no notable difference in the affinity (Fig. 2b). Our analyses do not exclude possible sequence preference or differential affinity for DNA with specific structure (e.g. bent, looped, or otherwise deformed)[28, 53].

Having established DNA binding by HupA_6xHis_, we examined the effect of replacing the arginine residues in the β-arm in the same assay. When HupA^QED^_6xHis_was incubated with all three tested DNA fragments, no shift was observed (Fig. 2a and b). This indicates that the introduction of the R55Q, R58E and R61D mutations successfully abolished binding of HupA to short dsDNA probes. We conclude that the arginine residues are crucial for the interaction with DNA and that the DNA-binding by HupA through the protruding β-arms is consistent with DNA binding by HU homologues from other organisms [35, 53, 57].

### Disruption of DNA binding does not affect oligomerization

HU proteins from various organisms have been found to form homo- or heterodimers [18, 19, 22, 53]. To determine the oligomeric state of *C. difficile* HupA protein, we performed size exclusion chromatography [60]. The elution profile of the purified protein was compared to molecular weight standards on a Superdex 75 HR 10/30 column. Wild-type HupA_6xHis_ protein exhibited a single clear peak with a partition coefficient (Kav) of 0.19 (Fig. 2c). These values correspond to an estimated molecular weight of a 38 kDa, suggesting a multimeric assembly of HupA_6xHis_ (theoretical molecular weight of monomer is 11 kDa). Similar to HupA_6xHis_, HupA^QED^_6xHis_ exhibits only one peak with a Kav of 0.20 and a calculated molecular weight of 37 kDa (Fig. 2c). Thus, mutation of the residues in the DNA-binding pocket of HupA did not interfere with the ability of HupA to form multimers in solution.

The calculated molecular weight for both proteins is higher than we would expect for a dimer (22 kDa), by analogy with HU proteins from other organisms. However, we cannot exclude the possibility that the conformation of the proteins affects the mobility in the size exclusion experiments. To better define the oligomeric state of HupA, we performed glutaraldehyde crosslinking experiments. HupA monomers cross-linked with glutaraldehyde were analyzed by western-blot analysis using anti-his antibodies. Upon addition of glutaraldehyde (0.0006% and 0.006%) we observed an additional signal around 23 kDa (Fig. 2d), consistent with a HupA dimer. No higher order oligomers were observed under the conditions tested. A similar picture was obtained for HupA^QED^_6xHis_ (Fig. 2d). Together, these experiments support the conclusion that HupA of *C. difficile* is a dimer in solution, similar to other described HU homologues, and that the ability to form dimers is independent of DNA-binding activity.

### *HupA self-interacts* in vivo

Above, we have shown that HupA of *C. difficile* forms dimers *in vitro.* We wanted to confirm that the protein also self-interacts *in vivo*. We developed a split-luciferase system to allow the assessment of protein-protein interactions in *C. difficile*. Our system is based on NanoBiT (Promega)[61] and our previously published codon-optimized variant of Nanoluc, sLuc^opt^ [62]. The system allows one to study protein-protein interactions *in vivo* in the native host, and thus present an advantage over heterologous systems. The large (LgBit) and small (SmBit) subunits of this system have been optimized for stability and minimal self-association by substitution of several amino acid residues [61]. When two proteins are tagged with these subunits and interact, the subunits come close enough to form an active luciferase enzyme that is able to generate a bright luminescent signal once substrate is added. We stepwise adapted our sLuc^opt^ reporter [62] by 1) removing the signal sequence (resulting in an intracellular luciferase, Luc^opt^), 2) introducing the mutations corresponding to the amino acid substitutions in NanoBiT (resulting in a full length luciferase in which SmBiT and LgBiT are fused, bitLuc^opt^) and finally, 3) the construction of a modular vector containing a polycistronic construct under the control of the anhydrotetracycline (ATc)-inducible promoter P_tet_ [63] (see Supplemental Methods).

To assess the ability of HupA to form multimers *in vivo*, we genetically fused HupA to the C-terminus of both SmBit and LgBit subunits and expressed them in *C. difficile* under the control of the ATc-inducible promoter. As controls, we assessed luciferase activity in strains that express full length luciferase (bitLuc^opt^) and combinations of HupA-fusions with or without the individual complementary subunit of the split luciferase (Fig. 3). Expression of the positive control bitLuc^opt^ results in a 2-log increase in luminescence signal after 1 hour of induction (1954024 ± 351395 LU/OD, Fig. 3). When both HupA-fusions are expressed from the same operon a similar increase in the luminescence signal is detected (264646 ± 122518 LU/OD at T1, Fig. 3). This signal is dependent on HupA being fused to both SmBit and LgBiT, as all negative controls demonstrate low levels of luminescence that do not significantly change upon induction (Fig. 3).

**Fig. 3.**
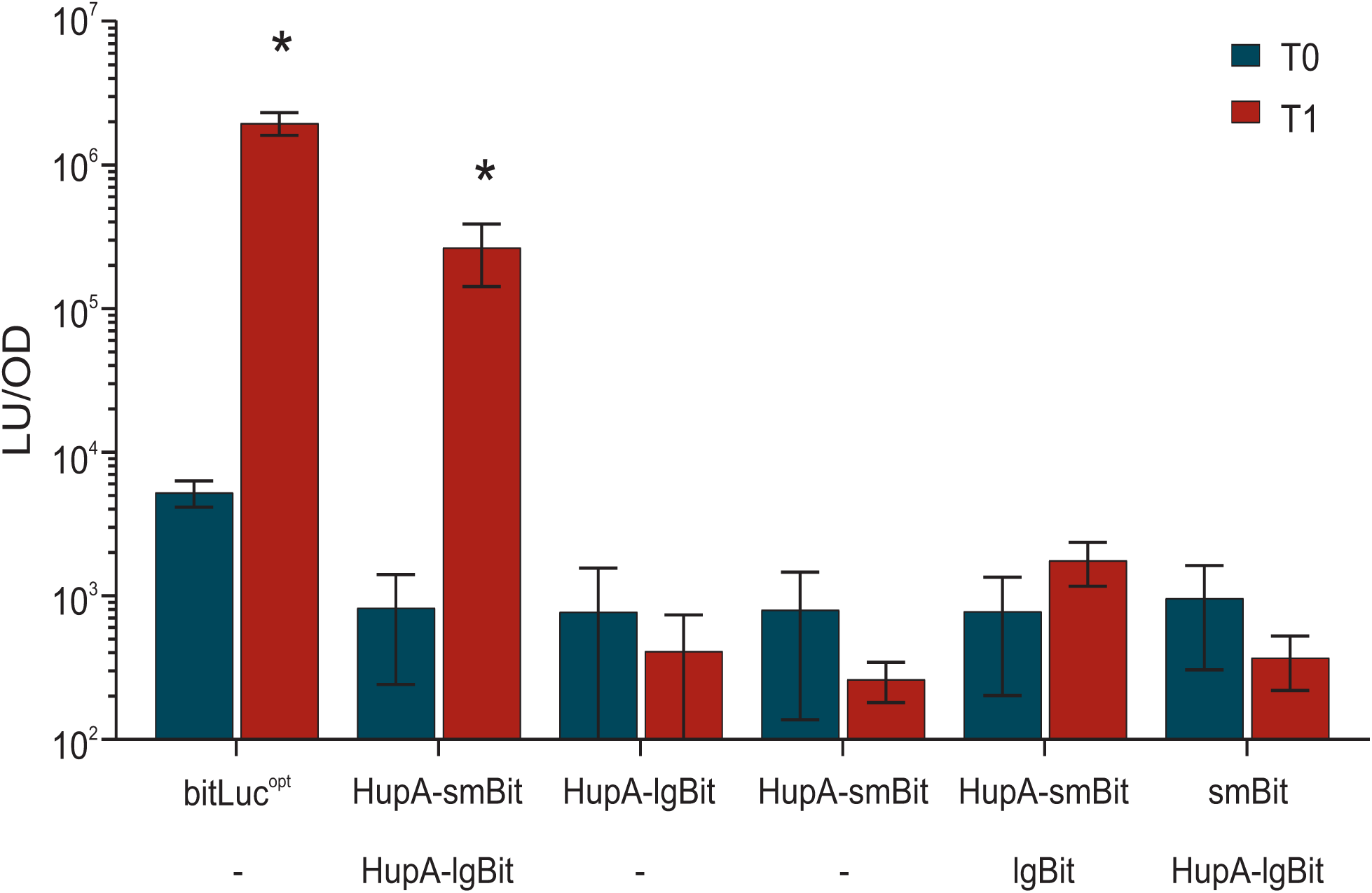
HupA demonstrates self-interaction in *C. difficile*. A split luciferase complementation assay was used to demonstrate interactions between HupA monomers *in vivo*. Cells were induced with 200 ng/mL anhydrotetracycline (ATc) for 60 min. Optical density-normalized luciferase activity (LU/OD) is shown right before induction (T0, blue bars) and after 1 hour of induction (T1, red bars). The averages of biological triplicate measurements are shown, with error bars indicating the standard deviation from the mean. Luciferase activity of strains AP182 (P_tet_-*bitluc*^*opt*^), AP122 (P_tet_-*hupA-smbit/hupA-lgbit*), AP152 (P_tet_-*hupA-lgbit*), AP153 (P_tet_-*hupA-smbit*), AP183 (P_tet_-*hupA-smbit-lgbit*) and AP184 (P_tet_-*smbit-hupA-lgbit*). A positive interaction was defined on the basis of the negative controls as a luciferase activity of >1000 LU/OD. No significant difference was detected at T0. AP122 and AP182 were significantly higher with p<0.0001 (*) by two-way ANOVA.

Our results indicate that HupA also self-interacts *in vivo.* However, we cannot exclude that the self-interaction is facilitated by other components of the cell (e.g. DNA or protein interaction partners).

### HupA overexpression leads to a condensed nucleoid

To determine if inducible expression of HupA leads to condensation of the chromosome in *C. difficile*, we introduced a plasmid carrying *hupA* under the ATc-inducible promoter P_tet_ into strain *630Δerm* [64]. This strain (AP106) also encodes the native *hupA* and induction of the plasmid-borne copy of the gene is expected to result in overproduction of HupA. AP106 cells were induced in exponential growth phase and imaged 1 hour after induction. In wild-type or non-induced AP106 cells nucleoids can be seen, after staining with DAPI stain, with a signal spread throughout most of the cytoplasm (Fig. 4a). In some cells a defined nucleoid is observed localized near the cell center (Fig. 4a). This heterogeneity in nucleoid morphology is likely a reflection of the asynchronous growth.

**Fig. 4.**
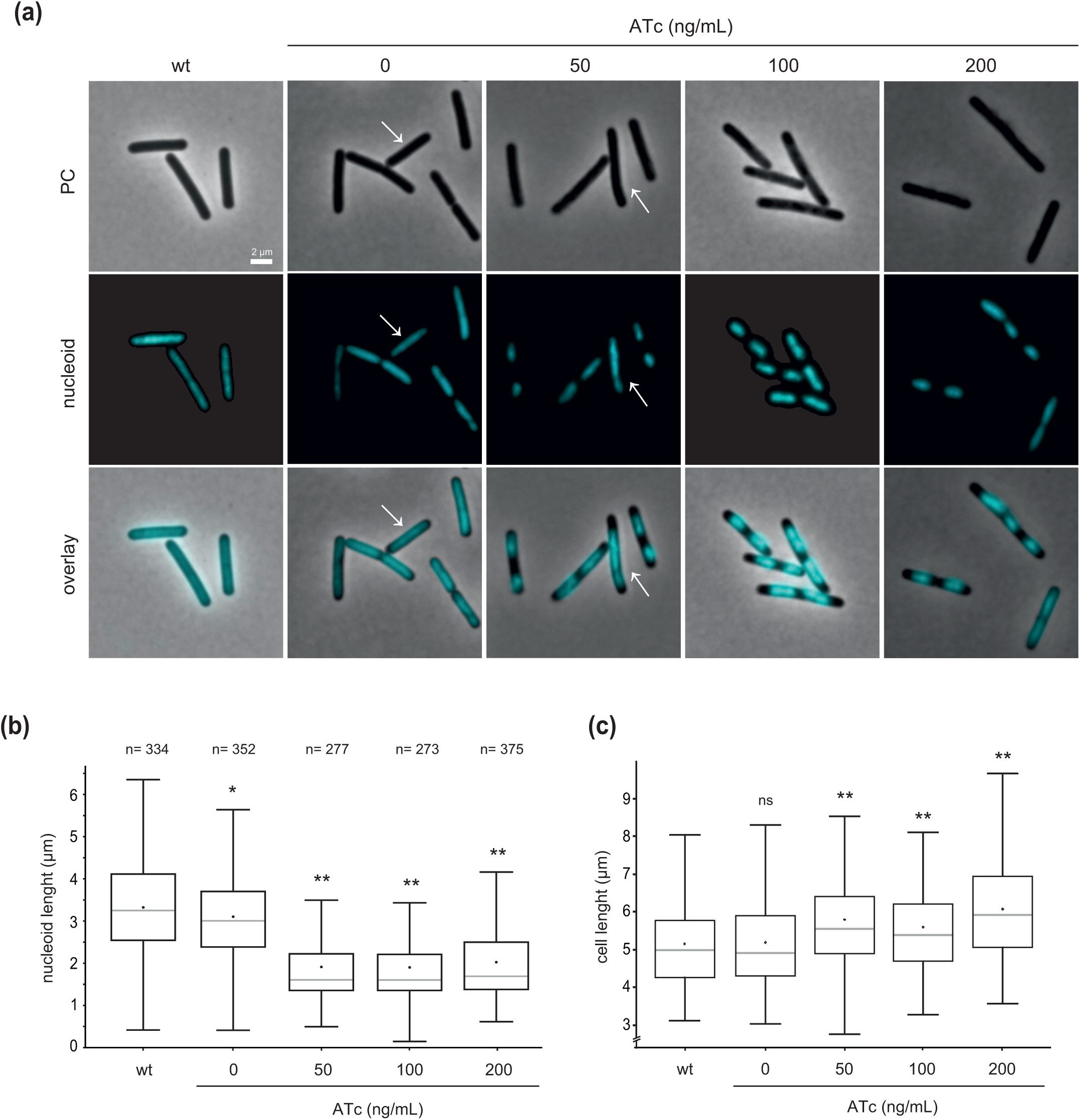
HupA overexpression leads to compaction of the nucleoid *in vivo*. (a) Fluorescence microscopy analysis of *C. difficile* 630Δ*erm* harboring the vector for anhydrotetracycline (ATc)-dependent overexpression of HupA (AP106). For HupA overexpression, cells were induced at mid-exponential growth in liquid medium with different ATc concentrations (50, 100 and 200 ng/mL) for 1h. *C. difficile* 630Δ*erm* and non-induced AP106 were used as controls. The cells were stained with DAPI for DNA visualization (nucleoid). The nucleoid was false colored in cyan for better contrast. Phase contrast (PC) and an overlay of both channels is shown. Because growth is asynchronous in these conditions, cells representing different cell cycles stages can be found. In the presence of ATc, the chromosome appears more compacted. White arrow indicates the cells with mid-cell nucleoid. Scale bar = 2 µm. (b) Boxplots of mean nucleoid length. Whiskers represent minimum and maximum nucleoid length observed. Black dots represent the mean and the grey lines represent the median values. Quantifications were performed using MicrobeJ from at least two biological replicates for each condition. n is the number of cells analyzed per condition. (c) Boxplots of mean cell length. Whiskers represent minimum and maximum cell length observed. Black dots represent the mean and the grey lines represent the median values. Quantifications were performed using MicrobeJ from at least two biological replicates for each condition. The same cells as analyzed for nucleoid length were used. * p<0,05 **p<0,0001 by one-way ANOVA compared to wild type (wt). ns = nonsignificant.

When HupA expression is not induced, the average nucleoid size is 3.10 ± 0.93 µm, similar to wild-type *C. difficile* 630Δ*erm* cells (3.32 ± 1.16 µm). Upon induction of HupA expression a significant decrease in size of the nucleoid is observed (Fig. 4a and b, white arrow). When cells are induced with 50, 100 or 200 ng/mL ATc the average nucleoid size was 1.91 ± 0.80 µm; 1.90 ± 0.82 µm and 2.02 ± 0.94 µm, respectively (Fig. 4b). No significant difference was detected between the strains induced with different ATc concentrations (Fig. 4b).

In wild-type *C. difficile* 630Δ*erm* cells the average cell length is 5.14 ± 1.07 μm, similar to non-induced AP106 cells (5.18 ± 1.09 μm, Fig. 4c). In the presence of increasing amounts of ATc a small but significant increase of cell length is observed after 1 hour induction. When cells are induced with 50, 100 or 200 ng/mL ATc the average cell length was 5.79 ± 1.29 µm; 5.58 ± 1.14 µm and 6.07 ± 1.37 µm, respectively (Fig. 4c). We did not observe an impairment of septum formation and localization (data not shown).

The decrease in the nucleoid size when HupA is overexpressed suggests that HupA can compact DNA *in vivo*. This observation is reminiscent of effects of HU overexpression reported for other organisms, like *B. subtilis* and *Mycobacterium tuberculosis* [10, 23].

### HupA co-localizes with the nucleoid

If HupA indeed is directly involved in condensing the nucleoid, it is expected that the protein co-localizes with the DNA. To test this, we imaged HupA protein and the nucleoid in live *C. difficile*. Here, we used the HaloTag protein (Promega)[65] for imaging the subcellular localization of HupA. Tags that become fluorescent after covalently labeling by small compounds, such as HaloTag, are proven to be useful for studies in bacteria and yeast [66–68]. Different from GFP, this type of tag does not require the presence of oxygen for maturation and should allow live-cell imaging in anaerobic bacteria.

We introduced a modular plasmid expressing HupA-HaloTag from the ATc-inducible promoter P_tet_ [63] into strain 630Δ*erm* [64], yielding strain RD16. Repeated attempts to create a construct that would allow us to integrate the fusion construct on the chromosome of *C. difficile* using allelic exchange failed, likely due to toxicity of the *hupA* upstream region in *E. coli* (cloning intermediate). For the visualization of HupA-HaloTag we used the Oregon green substrate, that emits at Em_max_ 520 nm. Although autofluorescence of *C. difficile* has been observed at wavelengths of 500-550 nm [69, 70] we observed limited to no green signal in the absence of the HaloTag (our unpublished observations and Fig. 5a,-ATc).

**Fig. 5.**
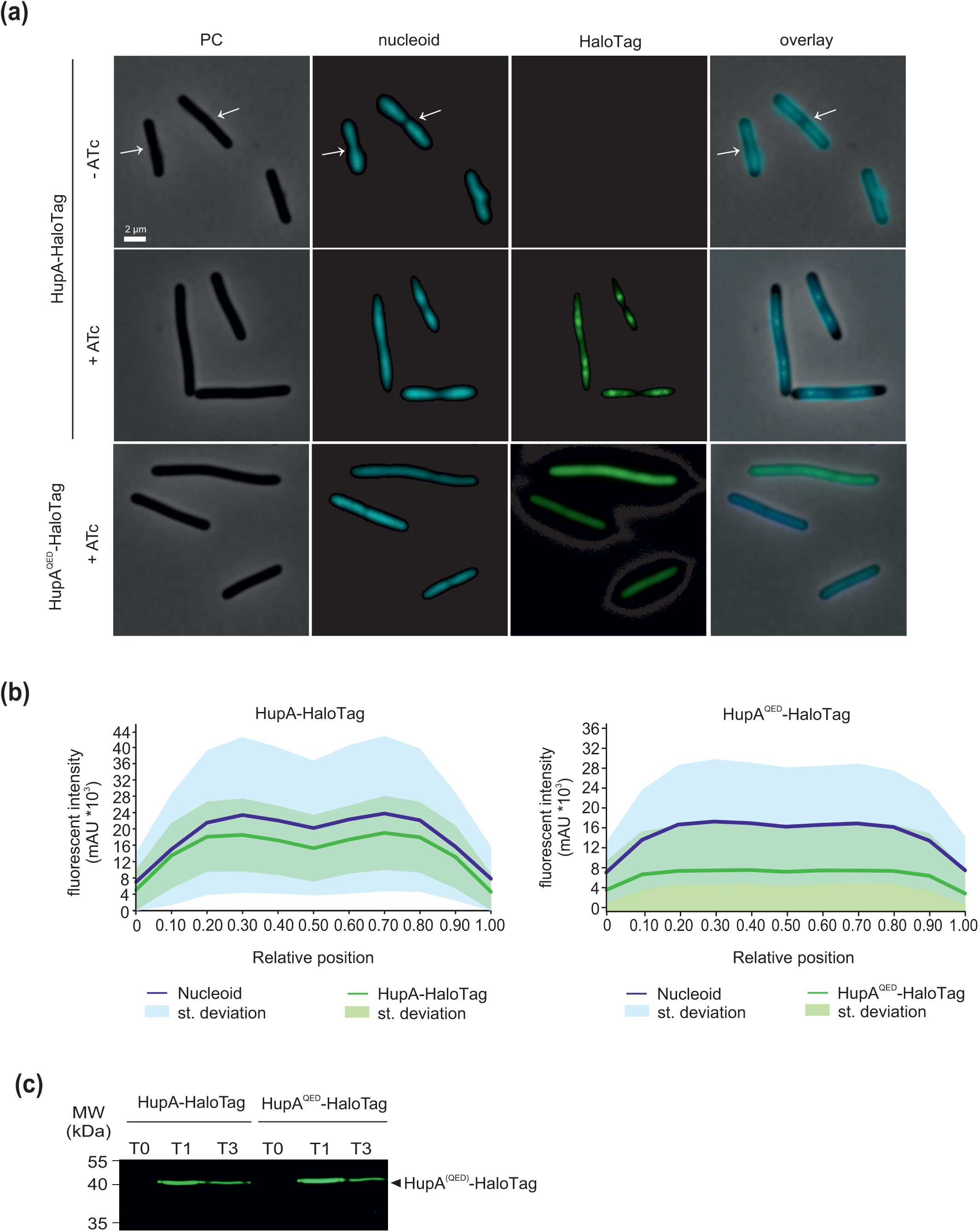
HupA co-localizes with the nucleoid. (a) Fluorescence microscopy analysis of *C. difficile* 630Δ*erm* harboring a vector for expression of HupA-HaloTag (RD16) or a vector for expression of HupA^QED^-HaloTag (AF239). For visualization of HupA-HaloTag and HupA^QED^-HaloTag, cells were induced at mid-exponential growth phase with 200 ng/mL anhydrotetracycline (ATc) for 1h and incubated with Oregon Green HaloTag substrate for 30 min. The cells were stained with DAPI stain to visualize DNA (nucleoid). The nucleoid was false colored in cyan for better contrast. As control non-induced RD16 is shown, but similar results were obtained for non-induced AF239. Phase contrast (PC) and an overlay of the channels is shown. Because growth is asynchronous under these conditions, cells representing different cell cycles stages can be found. In the presence of ATc the chromosome appears more compacted and HupA-HaloTag co-localizes with the nucleoid. Scale bar = 2 µm. (b) Average intensity profile scans for the nucleoid (DAPI, blue line) and HupA fusion protein (Oregon Green, green line) obtained from a MicrobeJ analysis from at least two biological replicates in each condition. 289 cells were analyzed for HupA-HaloTag and 331 cells were analyzed for HupA^QED^-HaloTag. Standard deviation of the mean is represented by the respective color shade. (c) In-gel fluorescent analysis of RD16 and AF239 samples before induction (T0), 1 and 3 hours after induction (T1, T3). Samples were incubated with Oregon Green substrate for 30 min and run on a 12% SDS-PAGE.

HupA-Halotag expression was induced in RD16 cells during exponential growth phase with 200 ng/mL ATc and cells were imaged after 1 hour of induction. In the absence of ATc, no green fluorescent signal is visible, and the nucleoid (stained with DAPI) appears extended (Fig. 5a). Upon HupA-HaloTag overexpression, the nucleoids are more defined and appear bilobed (Fig. 5a and b), similar to previous observations (Fig. 4a). The Oregon Green signal co-localizes with the nucleoid, located in the center of the cells, with a bilobed profile that mirrors the profile of the DAPI stain (Fig. 5a and b). This co-localization is observed for individual cells at different stages of the cell cycle and is independent of the number of nucleoids present (data not shown). The localization pattern of the *C. difficile* HupA resembles that of HU proteins described in other organisms [23, 71, 72] (Fig 5a). Expression levels of HupA-Halotag were confirmed by SDS-PAGE in-gel fluorescence of whole cell extracts, after incubation with Oregon Green (Fig. 5c).

ATc-induced RD16 cells exhibit a heterogeneous Oregon Green fluorescent signal. This has previously been observed with other fluorescent reporters in *C. difficile* [68–70, 73] and can likely be explained by both heterogeneous expression from inducible systems [74] and different stages of the cell cycle. For instance, the localization of cell division proteins, such as MldA or FtsZ is dependent on septum formation and thus dependent on cells undergoing cell division [69, 73].

We found that HupA^QED^_6xHis_ does not bind dsDNA in the electrophoretic mobility shift assay (Fig. 2b). We introduced the triple substitution in the HupA-HaloTag expression plasmid to determine its effect on localization of the protein in *C. difficile.* We found that the HupA^QED^-HaloTag protein was broadly distributed throughout the cell and that - different from ATc-induced RD16 cells (HupA-Halotag) - no compaction of the nucleoid occurred (Fig. 5a). The lack of compaction is not due to lower expression levels of HupA^QED^-Halotag compared to HupA-HaloTag, as induced levels of both proteins are similar (Fig. 5c).

The nucleoid morphology upon expression of HupA^QED^-HaloTag is similar to that observed in wild type 630Δ*erm* cells (Fig. 4a), suggesting that HupA^QED^ does not influence the activity of the native HupA *in vivo*. Though the mutated residues did not affect oligomerization (Fig. 2c and d) we considered the possibility that HupA^QED^ is unable to form heterodimers with native HupA. To evaluate whether HupA^QED^ and HupA can interact, we performed glutaraldehyde crosslinking and an *in vivo* complementation assay (Fig. S2). To allow for discrimination between momoners of wild type and mutant HupA in the crosslinking assay, we purified the HupA-HaloTag from *C. difficile* and incubated this protein with heterologously produced and purified HupA_6xHis_ or HupA^QED^. Upon crosslinking bands corresponding to dimers of the his-tagged (22 kDa) and the HaloTagged protein (96 kDa) are detectable (Fig. S2a), confirming our previous results (Fig. 2d). We also detect a signal corresponding to the molecular weight of a heterodimer with both HupA_6xhis_ and HupA^QED^_6xhis_ (56 kDa), suggesting that wild type and mutant protein can form heterodimers *in vitro* (Fig S2a). To analyze the *in vivo* behavior of these proteins, HupA^QED^ was expressed fused to SmBit and HupA to LgBit in the split luciferase complementation assay. In line with the crosslinking experiment, we observe luciferase reporter activity that is similar to that observed for AP122 (HupA-SmBiT/HupA-LgBiT). Thus, mutation of the arginine residues does not abolish the interaction with HupA *in vivo*. Nevertheless, it is conceivable that wild type homodimers are preferentially formed *in vivo* despite expression of HupA^QED^: the lack of DNA binding by HupA^QED^ could result in an effectively lower local concentration in the nucleoid than for wild type HupA.

Together, these results indicate that HupA co-localizes with the nucleoid, and that nucleoid compaction upon HupA overexpression is possibly dependent on its DNA-binding activity. We cannot exclude that the nucleoid compaction observed is an indirect result of HupA overexpression by influencing possible interaction with RNA and/or other proteins, or by altering transcription/translation [40, 75].

### *HupA compacts DNA* in vitro

To substantiate that the decrease in nucleoid size is directly attributable to the action of HupA, we sought to demonstrate a remodeling effect of HupA on DNA *in vitro*. We performed a ligase-mediated DNA cyclization assay. Previous work has established that a length smaller than 150 bp greatly reduces the possibility of the extremities of dsDNA fragments to meet. This makes the probability to ligate into closed rings less [76]. However, in the presence of DNA bending proteins exonuclease III (ExoIII)-resistant (thus closed) rings can be obtained [56, 76].

We tested the ability of HupA_6xHis_ to stimulate cyclization of a [γ-^32^P]-labeled 123-bp DNA fragment (Fig. 6a). The addition of T4 DNA-ligase alone results in multiple species, corresponding to ExoIII-sensitive linear multimers (Fig. 6a, lane 2 and 3). In the presence of HupA_6xHis_, however, an ExoIII-resistant band is visible (Fig. 6a, lanes 4 to 6). In the absence of ExoIII, the linear dimer is still clearly visible in the HupA-containing samples (Fig. 6a, last lane). We conclude that *C. difficile* HupA is able to bend the DNA, or otherwise stimulate cyclization by increasing flexibility and reducing the distance between the DNA fragment extremities, allowing the ring closure in the presence of ligase.

**Fig. 6.**
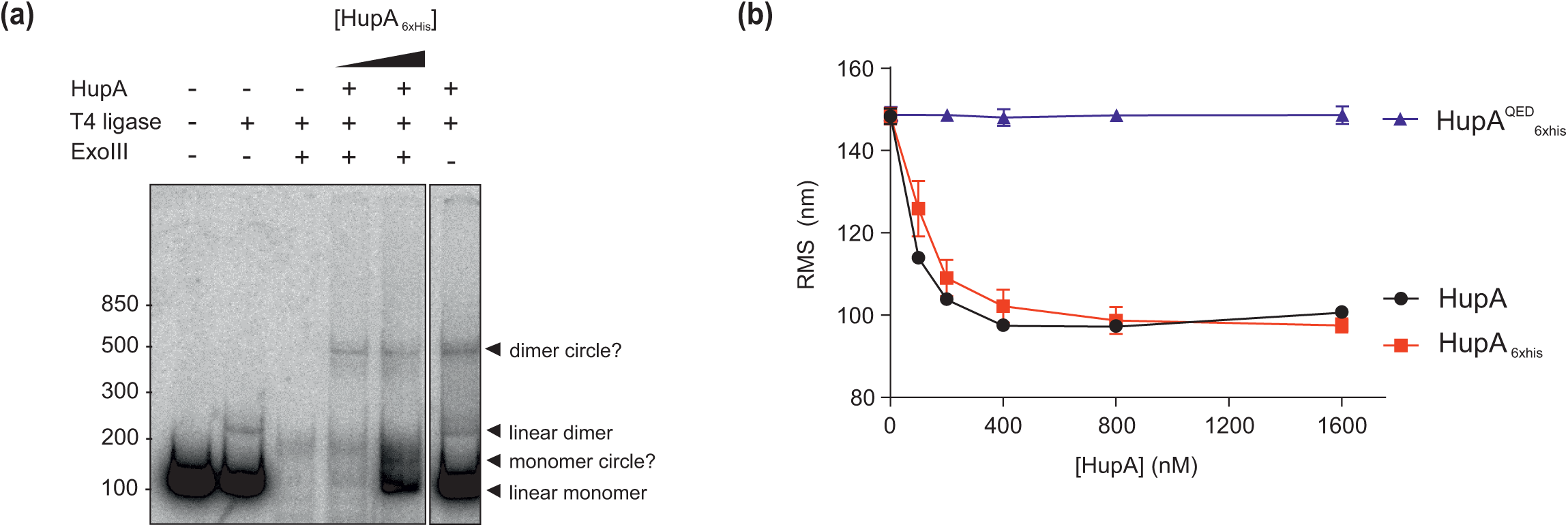
HupA alters the topology of DNA *in vitro*. (a) Ligase-mediated cyclization assay. A 119 bp [γ-^32^P]ATP-labelled dsDNA fragment was incubated in the presence of increasing concentrations of HupA_6xHis_ (1, 10 µM), exonuclease III and ligase, as indicated above the panel. The presence of ExoIII-resistant (i.e. circular) DNA fragments is observed when samples are incubated with HupA_6xHis_ (“circle”). (b) The effect of increasing concentrations of HupA (black circles), HupA_6xHis_ (red squares) and HupA^QED^_6xHis_ (blue triangles) on DNA conformation in tethered particle motion experiments. Root mean square (RMS; see Equation 1/Materials and methods) values as a function of protein concentration is shown. Increasing concentrations of HupA and HupA_6xHis_ lead to a decreased RMS, suggesting compaction of the DNA.

To more directly demonstrate remodeling of DNA by HupA, we performed tethered particle motion (TPM) experiments. TPM is a single molecule technique that provides a readout of the length and flexibility of a DNA tether (Supplemental Fig. S3)[77]. The binding of proteins to DNA alters its conformation, resulting in a change in RMS (Root Mean Square). If a protein bends DNA, makes DNA more flexible or more compact, the RMS is reduced compared to that of bare DNA, as represented in Supplemental Fig. S3 [77]. If a protein stiffens DNA, the RMS is expected to be larger than that of bare DNA [78].

We performed TPM experiments according to established methods [78] to determine the effects of HupA on DNA conformation at protein concentrations from 0 – 1600 nM (Fig. 6b). For this assay a non-tagged HupA was purified from *C. difficile* cells overexpressing HupA and compared to HupA_6xHis_ to assess potential subtle effects of the 6xhistidine-tag on the protein functionality. The experiments show that binding of both native HupA and HupA_6xHis_ to DNA reduces the RMS (Fig. 6b). The RMS of bare DNA is 148 ± 1.9 nm. In the presence of HupA at different concentrations (100, 200, 400 nM) the RMS decreases (113 ± 0.1 nm; 103 ± 0.7 nm and 97 ± 1.5 nm respectively). Even at higher concentrations of HupA (800, 1600 nM) the RMS is 97-100 nm. HupA^QED^_6xHis_ did not affect RMS even at high protein concentrations (Fig. 6b). The strongly reduced RMS of DNA bound by non-tagged HupA at 1600 nM suggests a more compacted conformation of DNA compared to that of bare DNA. The curves are overall highly similar for HupA and HupA_6xHis_ proteins; the small difference in the observed effects is attributed to interference of the tag and/or protein stability. The results obtained with the HupA^QED^_6xhis_protein indicate that DNA binding by HupA is crucial for compaction, as expected. We also tested the effect of the addition of a two-fold molar excess of HupA^QED^_6xhis_on DNA compaction in the presence of 400 nM HupA in a TPM experiment, but observed no significant difference (data not shown). This indicates that under these conditions HupA^QED^_6xhis_ does not remove DNA-bound HupA or affects its ability to remodel the DNA.

The effects of *C. difficile* HupA on DNA conformation observed by TPM indicate structural properties similar to those of *E. coli* HU, which was shown to compact DNA by bending at low protein coverage [26, 79, 80]. However, in contrast to *E. coli* [26], there is no clear stiffening of the DNA tether at high concentrations of protein in our assay, suggesting that there is lower or reduced dimer-dimer interaction in our experimental condition. Bending of DNA by HU proteins has also been shown for other organisms. Interestingly, in *B. burgdorferi* [55] and *Anabaena* [35] it was shown that bending is influenced by interaction of the DNA with a positively charged lateral surface, although the main interaction region with the DNA is through the β-arms. *C. difficile* HupA demonstrates an electrostatic surface potential compatible with such a mechanism (Fig 1C). It will be of interest to determine if and which residues in this region contribute to the bending of the DNA.

### Overexpression of HupA decreases cell viability

The condensation of the nucleoid and the slight increase of cell length during the time course of our microscopy experiments (Fig. 4b and c) could indicate that overexpression of HupA interferes with crucial cellular processes such as DNA replication. We therefore determined the long term effect of HupA overexpression on cell viability in a spot-assay (Fig. 7). In the absence of inducer, *C. difficile* strains harboring inducible *hupA* genes grow as well as the vector control (AP34), with colonies visible at the 10^-5^ dilution. However, when induced with 200 ng/ml ATc viability is markedly reduced for strains overexpressing HupA (5-log; AP106), HupA-HaloTag (4-log; RD16) and HupA^QED^-HaloTag (1 to 2-log; AF239) compared to the vector control. These effects are not due to a direct inhibitory effect of ATc alone, as the viability of AP34 is similar under both conditions.

**Fig. 7.**
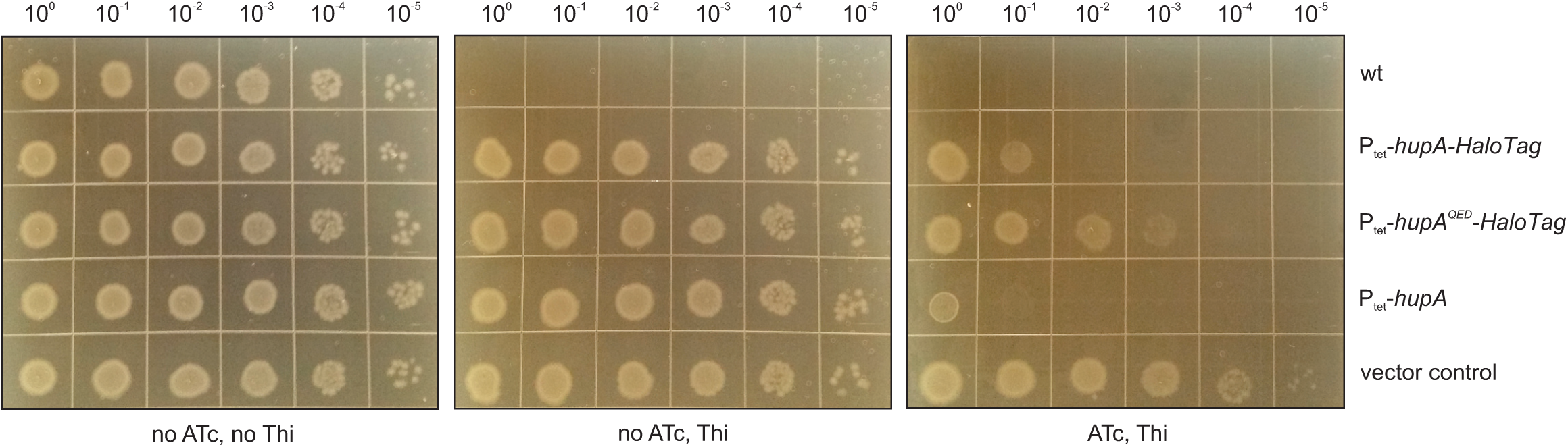
Strain viability under conditions of HupA overexpression. Spot assay of serially diluted *C. difficile* strains 630Δ*erm,* RD16 (P_tet_-*hupA-HaloTag*), AF239 (P_tet_-*hupA*^*QED*^*-Halotag*), AP106 (P_tet_-*hupA*) and AP34 (P_tet_-*sluc*^*opt*^). The left panel shows growth on medium with only *C. difficile* selective supplement (CDSS), the middle panel shows growth on medium with CDSS and thiamphenicol (Thi) and the right panel shows growth on medium with CDSS, Thi and 200 ng/mL anhydrotetracyclin (ATc) after 24 hours at 37ºC. The results were verified by four independent spot assays and a typical image is shown. Overexpression of HupA strongly reduces cell viability.

We consistently observed a 1-log difference in cell viability between cells expressing HupA versus HupA-HaloTag (Fig. 7). This difference could be the result of slight interference of the HaloTag with HupA function, as also observed for the 6xhistagged protein in the TPM experiments (Fig. 6b). Considering that HupA^QED^ does not appear to bind or compact DNA (Figs. 2, 5 and 6), the moderate reduction in cell viability compared to the vector control could be due to a dominant negative effect: the formation of heterodimers, consistent with our analysis (Fig. S2), could prevent a fraction of wild type HupA performing its essential function.

Overall, these results are consistent with an important role of HupA in chromosome dynamics.

## Conclusions

In this work, we present the first characterization of a bacterial chromatin protein in *C. difficile*. HupA is a member of the HU family of proteins and is capable of binding DNA and does so without an obvious difference in affinity as a result of the G+C content. DNA binding is dependent on the residues R55, R58 and R61 that are located in the predicted β-arm of the protein. These observations in combination with the predicted structure suggest a conserved mode of DNA binding, although the role of other regions of the protein in DNA binding is still poorly understood. HupA is present as a dimer in solution and disruption of the residues of the DNA binding domain did not affect the oligomeric state of HupA.

In *C. difficile* we co-localized HupA with the nucleoid and demonstrated that overexpression of HupA leads to nucleoid compaction and impairs *C. difficile* viability. In line with these observations, HupA stimulates the cyclization of a short dsDNA fragment and compacts DNA *in vitro*.

We also developed a new complementation assay for the detection of protein-protein interactions in *C. difficile,* complementing the available tools for this organism, and confirmed that HupA self-interacts *in vivo*. Additionally, to our knowledge, our study is the first to describe the use of the fluorescent tag HaloTag for imaging the subcellular localization of proteins in live *C. difficile* cells.

In sum, HupA of *C. difficile* is an essential bacterial chromatin protein required for nucleoid (re)modelling. HupA binding induces bending or increases the flexibility of the DNA, resulting in compaction. The precise role of HupA in chromosome dynamics *in vivo* remains to be determined. In *E. coli* conformational changes resulting from HU proteins enhance contacts between distant sequences in the chromosome [81]. In *Caulobacter*, HU proteins promote contacts between sequences in more close proximity [82]. These differences demonstrate that HU proteins despite high sequence similarity may act differently as a function of *in vivo* context and that further research into the role of HupA in *C. difficile* physiology is needed.

## Methods

### Sequence Alignments and Structural Modelling

Multiple sequence alignment of amino acid sequences was performed with Clustal Omega [83]. The sequences of HU proteins identified in *C. difficile* 630Δ*erm* (Q180Z4), *E. coli* (P0ACF0 and P0ACF4), *Bacillus subtilis* (A3F3E2), *Geobacillus stearothermophilus* (P0A3H0), *Bacillus anthracis* (Q81WV7), *Staphylococcus aureus* (Q99U17), *Salmonella typhimurium* (P0A1R8), *Streptococcus pneumoniae* (AAK75224), *S. mutans* (Q9XB21), *M. tuberculosis* (P9WMK7), *Thermotoga maritima* (P36206) and *Anabaena sp.* (P05514), were selected for alignment. Amino acid sequences were retrieved from the Uniprot database.

Homology modelling was performed using PHYRE2 (http://www.sbg.bio.ic.ac.uk/phyre2, [51] and SWISS-MODEL [52] using default settings. For SWISS-MODEL, PDB 4QJN was used as a template. Selection of the template was based on PHYRE2 results, sequence identity (59,55%) and best QSQE (0,80) and GMQE (0,81). Graphical representations and mutation analysis were performed with the PyMOL Molecular Graphics System, Version 1.76.6. Schrödinger, LLC. For electrostatics calculations APBS (Adaptive Poisson-Boltzmann Solver) and PDB2PQR software packages were used [84]. Default settings were used.

### Strains and growth conditions

*E. coli* strains were cultured in Luria Bertani broth (LB, Affymetrix) supplemented with chloramphenicol at 15 µg/mL or 50 µg/mL kanamycin when appropriate, grown aerobically at 37°C. Plasmids (Table 1) were maintained in *E. coli* strain DH5α. Plasmids were transformed using standard procedures [85]. *E. coli* strain Rosetta (DE3) (Novagen) was used for protein expression and *E. coli* CA434 for plasmid conjugation [86] with *C. difficile* strain 630Δ*erm* [64, 87].

**Table 1.**
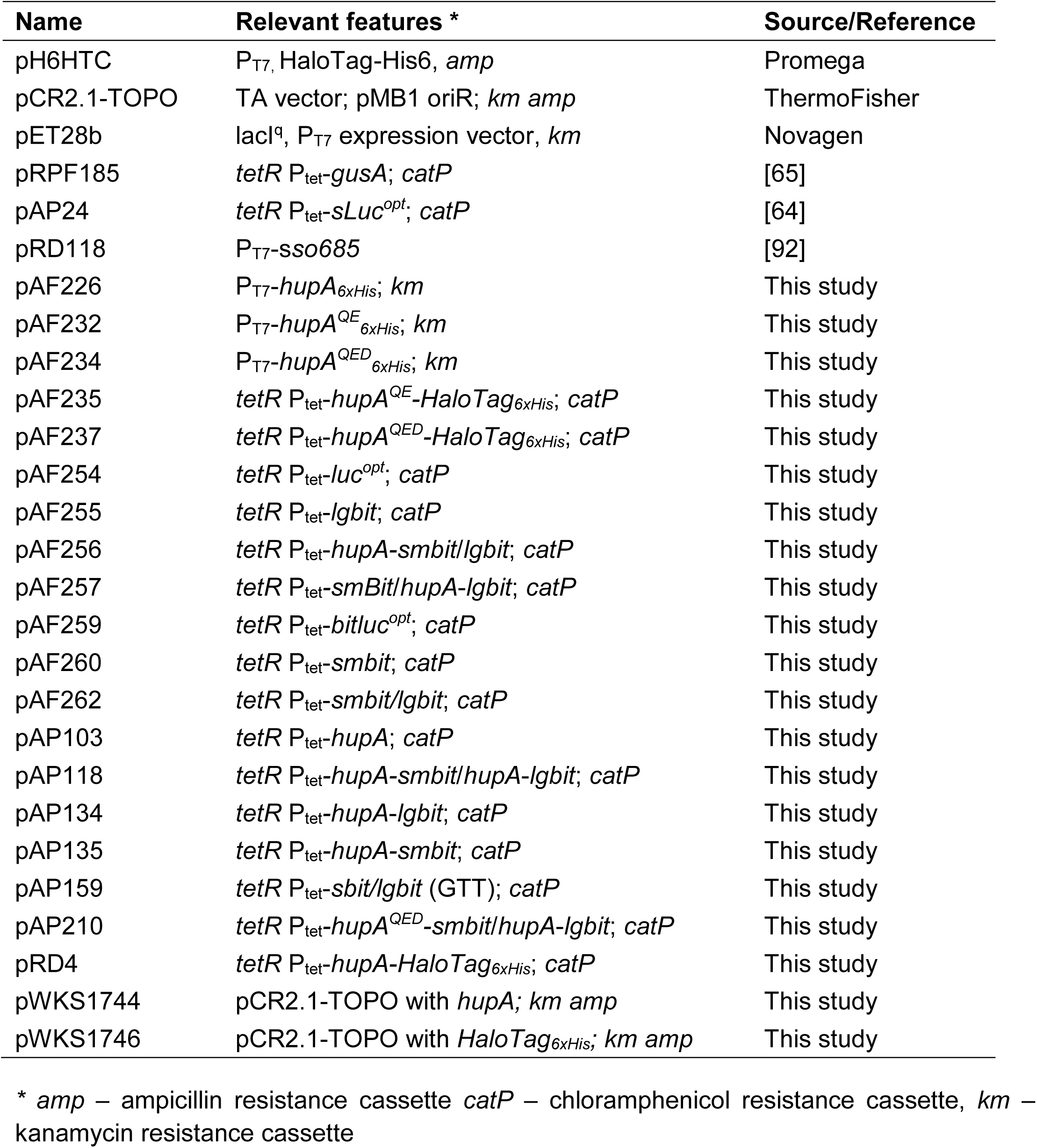
Plasmids used in this study.

*C. difficile* strains were cultured in Brain Heart Infusion broth (BHI, Oxoid), with 0,5 % w/v yeast extract (Sigma-Aldrich), supplemented with 15 µg/mL thiamphenicol and *Clostridium difficile* Selective Supplement (CDSS; Oxoid) when necessary. *C. difficile* strains were grown anaerobically in a Don Whitley VA-1000 workstation or a Baker Ruskinn Concept 1000 workstation with an atmosphere of 10% H_2_, 10% CO_2_ and 80% N_2_.

The growth was followed by optical density reading at 600 nm. All the *C. difficile* strains are described in Table 2.

**Table 2.** *C. difficile* strains used in this study.

| **Name** | **Relevant Genotype/Phenotype\*** | **Source/Reference** |
| --- | --- | --- |
| AP6 | *C. difficile* 630Δ*erm*; Erm<sup>S</sup> | [66, 87] |
| WKS1588 | 630Δ*erm* pRPF185; Thia<sup>R</sup> | This study |
| RD16 | 630Δ*erm* pRD4; Thia<sup>R</sup> | This study |
| AF239 | 630Δ*erm* pAF237; Thia<sup>R</sup> | This study |
| AP34 | 630Δ*erm* pAP24; Thia<sup>R</sup> | [64] |
| AP106 | 630Δ*erm* pAP103; Thia<sup>R</sup> | This study |
| AP122 | 630Δ*erm* pAP118; Thia<sup>R</sup> | This study |
| AP152 | 630Δ*erm* pAP134; Thia<sup>R</sup> | This study |
| AP153 | 630Δ*erm* pAP135; Thia<sup>R</sup> | This study |
| AP181 | 630Δ*erm* pAF254; Thia<sup>R</sup> | This study |
| AP182 | 630Δ*erm* pAF259; Thia<sup>R</sup> | This study |
| AP183 | 630Δ*erm* pAF256; Thia<sup>R</sup> | This study |
| AP184 | 630Δ*erm* pAF257; Thia<sup>R</sup> | This study |
| AP199 | 630Δ*erm* pAF255; Thia<sup>R</sup> | This study |
| AP201 | 630Δ*erm* pAF260; Thia<sup>R</sup> | This study |
| AP202 | 630Δ*erm* pAF262; Thia<sup>R</sup> | This study |
| AP212 | 630Δ*erm* pAP210; Thia<sup>R</sup> | This study |

\* Erm<sup>S</sup> – Erythromycin sensitive, Thia<sup>R</sup> – Thiamphenicol resistant

### Construction of the E. coli expression vectors

All oligonucleotides and plasmids from this study are listed in Tables 1 and 3.

**Table 3.**
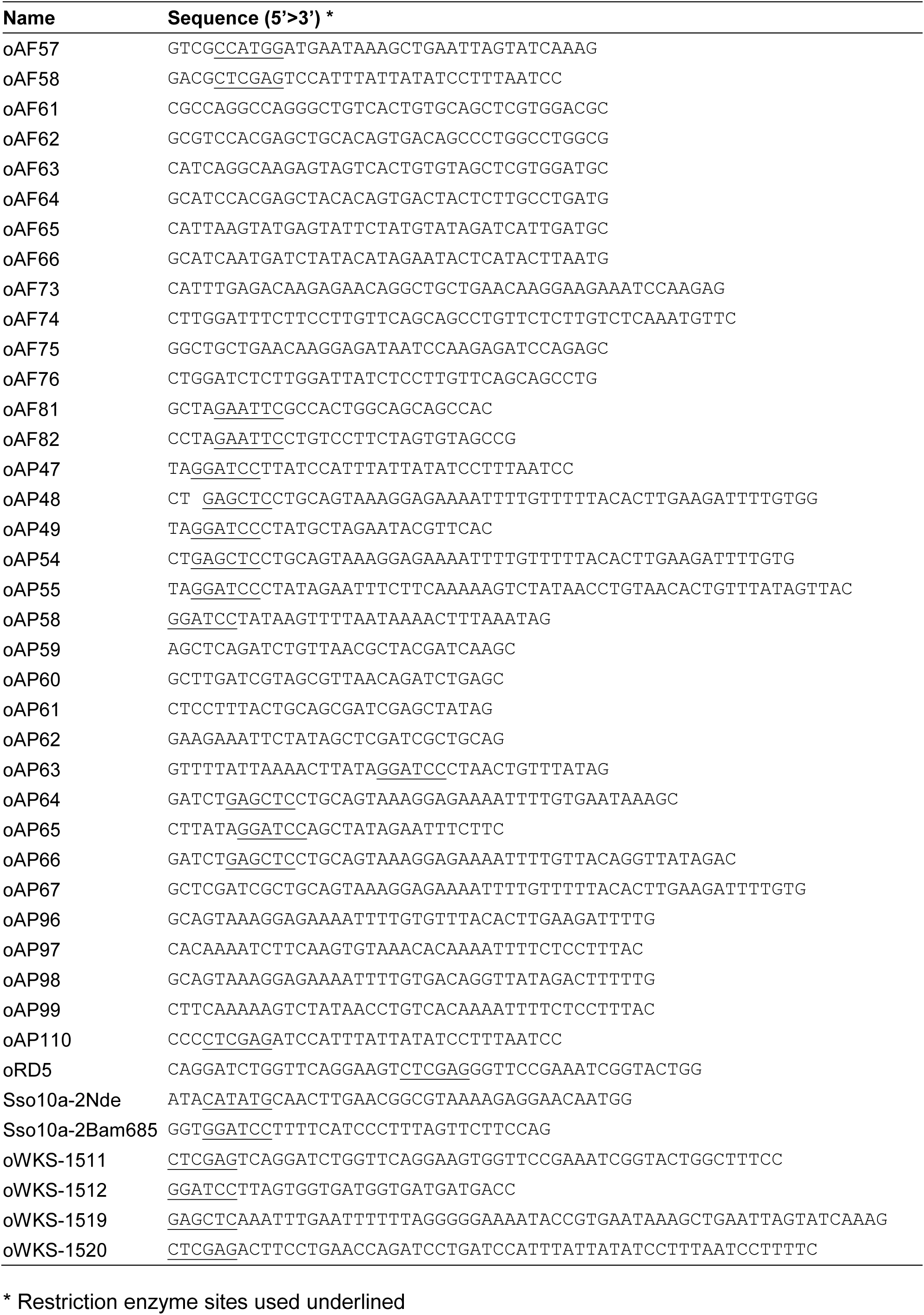
Oligonucleotides used in this study.

To construct an expression vector for HupA_6xHis_, the *hupA* gene (CD3496 from *C. difficile* 630 GenBank accession no. NC_009089.1) was amplified by PCR from *C. difficile* 630Δ*erm* genomic DNA using primers oAF57 and oAF58 (Table 3). The product was inserted into the *Nco*I-*Xho*I digested pET28b vector (Table 1) placing it under control of the T7 promoter, yielding plasmid pAF226.

To generate the HupA triple mutant (HupA^QED^_6xHis_) site-directed mutagenesis was used according to the QuikChange protocol (Stratagene). Initially the arginine at position 55 and at position 58 were simultaneously substituted for glutamine (R55Q) and glutamic acid (R58E) respectively, using primers oAF73/oAF74 (Table 3), resulting in pAF232 (Table 1). The arginine at position 61 was subsequently substituted for aspartic acid (R61D) using primer pair oAF75/oAF76 (Table 3) and pAF232 as a template, yielding pAF234 (Table 1). All the constructs were confirmed by Sanger sequencing.

### Construction of the C. difficile expression vectors

To overexpress non-tagged HupA the *hupA* gene was amplified by PCR from *C. difficile 630Δerm* genomic DNA using primers oWKS-1519 and oAP47 (Table 3) and cloned into SacI-BamHI digested pRPF185 vector [63], placing it under control of the ATc-inducible promoter P_tet_, yielding vector pAP103 (Table 1).

For microscopy experiments HaloTagged protein (Promega) was used. The *halotag* gene was amplified from vector pH6HTC (Promega, GenBank Accession no. JN874647) with primers oWKS-1511/oWKS-1512 and inserted into pCR2.1-TOPO according to the instructions of the manufacturer (ThermoFisher), yielding vector pWKS1746 (Table 1). This primer combination also introduces a 6xHis-tag at the C-terminus of the HaloTag. The *hupA* gene was amplified with primers oWKS-1519/oWKS-1520 (Table 3) and inserted into vector pCR2.1-TOPO according to the instructions of the manufacturer (ThermoFisher), generating vector pWKS1744 (Table 1). The primers introduces the *cwp2* ribosomal binding site upstream and a short DNA sequence encoding a GS-linker downstream (SGSGSGS) of the *hupA* open reading frame. To generate the expression construct for HupA-Halotag the open reading frame encoding the HaloTag_6xHis_ protein was amplified from pWKS1746 using primers oRD5/oWKS-1512 (Table 3). The *hupA* gene was amplified from pWKS1744 with primers oWKS-1519/oWKS-1520 (Table 3). Gene fusions were made by overlapping PCR using the PCR amplified fragments encoding HupA and Halotag proteins as templates with primers oWKS-1519 and oWKS-1512 (Table 3). The fragment was cloned into SacI-BamHI digested pRPF185 [63], placing it under control of the ATc inducible promoter P_tet_, yielding vector pRD4 (Table 1).

To generate the HupA triple mutant fused to the Halotag (HupA^QED^-Halotag) site-directed mutagenesis was used, according to the QuikChange protocol (Stratagene). The arginines at position 55 and at position 58 were substituted to glutamine (R55Q) and glutamic acid (R58E), using primers oAF73/oAF74 (Table 3) and pRD4 as template, resulting in pAF235 (Table 1). The arginine at position 61 was subsequently substituted to aspartic acid (R61D), using pAF235 as template and primers oAF75/oAF76 (Table 3), yielding pAF237 (Table 1). All the constructs were confirmed by Sanger sequencing.

### *Construction of the bitLuc*^opt^ *expression vectors*

The bitLuc^opt^ complementation assay for *C. difficile* described in this study is based on NanoBiT (Promega) [61] and the codon optimized sequence of sLuc^opt^ [62]. Details of its construction can be found in Supplemental Material.

Gene synthesis was performed by Integrated DNA Technologies, Inc. (IDT). Fragments were amplified by PCR from synthesized dsDNA, assembled by Gibson assembly [88] and cloned into *Sac*I/*BamH*I digested pRPF185 [63], placing them under control of the ATc-inducible promoter P_tet_. As controls a non-secreted luciferase (Luc^opt^; pAF254) and a luciferase with the NanoBiT aminoacid substitutions (Promega)[61](bitLuc^opt^; pAF259) were constructed. We also constructed vectors expressing only the SmBiT and LgBiT domains, alone (pAF260 and pAF255) or in combination (pAF262), as controls.

To assay for a possible interaction between HupA monomers, vectors were constructed that encode HupA-SmBiT/HupA-LgBiT (pAP118), HupA^QED^-SmBiT/HupA-LgBiT (pAP210), HupA-SmBiT/LgBiT (pAF256), SmBiT/HupA-LgBiT (pAF257). DNA sequences of the cloned DNA fragments in all recombinant plasmids were verified by Sanger sequencing.

Note that all our constructs use the HupA start codon (GTG) rather than ATG; a minimal set of vectors necessary to perform the *C. difficile* complementation assay (pAP118, pAF256, pAF257 and pAF258) is available from Addgene (105494-105497) for the *C. difficile* research community.

### *Overproduction and purification of* HupA^(QED)^_6xhis_ and HupA-HaloTag

Overexpression of HupA_6xHis_ and HupA^QED^_6xHis_ was carried out in *E. coli* Rosetta (DE3) strains (Novagen) harbouring the *E. coli* expression plasmids pAF226 and pAF234, respectively. Cells were grown in LB and induced with 1mM isopropyl-β-D-1-thiogalactopyranoside (IPTG) at an optical density (OD_600_) of 0,6 for 3 hours. The cells were collected by centrifugation at 4°C and stored at −80°C.

Overexpression of HupA-HaloTag (which also includes a 6xhistag) was carried out in *C. difficile* strains RD16. Cells were grown until OD_600_ 0.4-0.5 and induced with 200 ng/mL ATc for 1 hour. Cells were collected by centrifugation at 4°C and stored at −80°C.

Pellets were suspended in lysis buffer (50 mM NaH_2_PO_4_ (pH 8.0), 300 mM NaCl, 10 mM imidazole, 5 mM β mercaptoethanol, 0.1% NP40 and Complete protease inhibitor cocktail (CPIC, Roche Applied Science). Cells were lysed by the addition of 1 mg/ml lysozyme and sonication. The crude lysate was clarified by centrifugation at 13000 g at 4°C for 20 min. The supernatant containing recombinant proteins was collected and purification was performed with TALON Superflow resin (GE Healthcare) according to the manufacturer’s instructions. Proteins were stored at −80°C in 50 mM NaH_2_PO_4_ (pH 8.0), 300 mM NaCl and 12% glycerol.

### Overproduction and purification of non-tagged HupA

Overexpression of HupA was carried out in *C. difficile* strain AP106 that carries the plasmid encoding HupA under the ATc-inducible promoter P_tet_. Cells were grown until OD_600_ 0.4-0.5 and induced with 200 ng/mL ATc for 3 hours. Cells were collected by centrifugation at 4°C.

Pellets were resuspended in HB buffer (25 mM Tris (pH 8.0), 0.1 mM EDTA, 5 mM β mercaptoethanol, 10% glycerol and CPIC). Cells were lysed by French Press and phenylmethylsulfonyl fluoride was added to 0.1 mM. Separation of the soluble fraction was performed by centrifugation at 13000g at 4°C for 20 min. Purification of the protein from the soluble fraction was done on a 1 mL HiTrap SP (GE Healthcare) according to manufacturer instructions. The protein was collected in HB buffer supplemented with 300 mM NaCl. Fractions containing the HupA protein were pooled together and applied to a 1 mL Heparin Column (GE Healthcare) according to manufacturer’s instructions. Column washes were performed with a 500 mM – 800 mM NaCl gradient in HB buffer. Proteins were eluted in HB buffer supplemented with 1 M NaCl and stored in 10% glycerol at −80°C.

### DNA labelling and electrophoretic mobility shift Assay (EMSA)

For the gel shift assays double stranded oligonucleotides with different GC contents were used. Oligonucleotides oAF61/oAF62 have a 71,1% G+C-content, oAF63/oAF64 have a 52,6% G+C-content and oAF65/oAF66 have a 28,9% G+C-content. The oligonucleotides were labelled with [γ-^32^P]ATP and T4 polynucleotide kinase (PNK) (Invitrogen) according to the PNK-manufacturer’s instructions. The fragments were purified with a Biospin P-30 Tris column (BioRad). Oligonucleotides with same G+C content were annealed by incubating them at 95°C for 10 min, followed by ramping to room temperature.

Gel shift assays were performed with increasing concentrations (0,25 −2 µM) of HupA_6xHis_ or HupA^QED^_6xHis_ in a buffer containing 20 mM Tris pH 8.0; 50 mM NaCl; 12 mM MgCl_2_; 2.5 mM ATP; 2 mM DTT; 10% glycerol and 2.4 nM [γ^-32^P]ATP-labelled oligonucleotides. Proteins were incubated with the oligonucleotide substrate for 20 min at room temperature prior to separation. Reactions were analyzed in 8% native polyacrylamide gels in cold 0,5X TBE buffer supplemented with 10 mM MgCl_2_. After electrophoresis gels were dried under vacuum and protein-DNA complexes were visualized by phosphorimaging (Typhoon 9410 scanner; GE Healthcare). Analysis was performed with Quantity-One software (BioRad).

### Size-exclusion chromatography

Size-exclusion experiments were performed on an Äkta pure 25L1 instrument (GE Healthcare). 200 μL of HupA_6xHis_ and HupA^QED^_6xHis_ was applied at a concentration of *∼*100 µM, to a Superdex 75 HR 10/30 column (GE Healthcare), in buffer containing 50 mM NaH_2_PO_4_ (pH 8.0), 300 mM NaCl and 12% glycerol. UV detection was done at 280 nm. Lower concentrations of HupA were not possible to analyses due to the lack of signal. HupA protein only contains 3 aromatic residues, and lacks His, Trp, Tyr or Cys to allow detection by absorbance at 280 nm. The column was calibrated with a mixture of proteins of known molecular weights (Mw): Conalbumin (75 kDa), Ovalbumin (44 kDa), Carbonic anhydrase (29 kDa), Ribonuclease A (13,7 kDa), and Aprotinin (6,5 kDa). Molecular weight of the HupA proteins was estimated according to the equation MW=10(Kav-b)/m where m and b correspond to the slope and the linear coefficient of the plot of the logarithm of the MW as a function of the Kav. The Kav is given by the equation Kav=(V_e_-V_0_)/(V_t_-V_0_) [89], where V_e_ is the elution volume for a given concentration of protein, V_0_ is the void volume (corresponding to the elution volume of thyroglobulin), and V_t_ is the total column volume (estimated from the elution volume of a 4% acetone solution).

### Glutaraldehyde cross-linking Assay

100 ng HupA protein was incubated with different concentrations of glutaraldehyde (0 0,006%) for 30 min at room temperature. Reactions were quenched with 10 mM Tris. The samples were loaded on a 6.5% SDS-PAGE gel and analysed by western-blotting. The membrane was probed with a mouse anti-His antibody (Thermo Fisher) 1:3000 in phosphate buffered saline (PBS; 137 mM NaCl, 2.7 mM KCl, 4.3 mM Na_2_HPO_4_, 1.47 mM KH_2_PO_4_) with 0.05% Tween-20 and 5% w/v Milk (Campina), a secondary anti-mouse HRP antibody 1:3000 and Pierce ECL2 Western blotting substrate (Thermo Scientific). A Typhoon 9410 scanner (GE Healthcare) was used to record the chemiluminescent signal.

### Split luciferase (bitLuc^opt^) Assay

For the *C. difficile* complementation assay, cells were grown until OD_600_ 0.3-0.4 and induced with 200 ng/mL anhydrotetracline for 60 min. To measure luciferase activity 20 µL NanoGlo Luciferase (Promega N1110) was added to 100 µL of culture sample. Measurements were performed in triplicate in a 96-well white F-bottom plate according to manufacturer’s instructions. Luciferase activity was determined using a GloMax instrument (Promega) for 0.1 s. Data was normalized to culture optical density measured at 600 nm (OD_600_). Statistical analysis was performed with Prism 7 (GraphPad, Inc, La Jolla, CA) by two-way ANOVA.

### Ligase-mediated cyclization assay

A 119 bp DNA fragment was amplified by PCR amplification with primers oAF81/oAF82, using pRPF185 plasmid as a template. The PCR fragment was digested with *EcoR*I and 5’end labelled with [γ3^2^P] ATP using T4 polynucloetide kinase (Invitrogen) according to the manufacturer’s instructions. Free ATP was removed with a Biospin P-30 Tris column (BioRad).

The labelled DNA fragment (*∼*0.5 nM) waspAF235 incubated with different concentrations of HupA for 30 min on 30°C in 50 mM Tris-HCl, pH 7.8, 10 mM MgCl_2_, 10 mM DTT, and 0.5 mM ATP in a total volume of 10 µl. 1 Unit of T4 ligase was added and incubated for 1 h at 30°C followed by inactivation for 15 minutes at 65°C. When appropriate, samples were treated with 100 U of Exonuclease III (Promega) at 37°C for 30 minutes. Enzyme inactivation was performed by incubating the samples for 15 minutes at 65°C. Before electrophoresis the samples were digested with 2 µg proteinase K and 0.2% SDS at 37°C for 30 minutes. Samples were applied to a pre-run 7% polyacrylamide gel in 0.5X TBE buffer with 2% glycerol and run at 100V for 85 min. After electrophoresis the gel was vacuum-dried and analysed by phosphor imaging. Analysis was performed with Quantity-One software (BioRad).

### Fluorescence microscopy

The sample preparation for fluorescence microscopy was carried out under anaerobic conditions. *C. difficile* strains were cultured in BHI/YE, and when appropriate induced with different ATc concentrations (50, 100 and 200 ng/mL) for 1 hour at an OD_600_ of 0.3-0.4. When required, cells where incubated with 150 nM Oregon Green substrate for HaloTag (Promega) for 30 min. 1 mL culture was collected and washed with pre-reduced PBS. Cells were incubated with 1 μM DAPI (Roth) when necessary. Cells were spotted on 1.5% agarose patches with 1 μL of ProLong Gold antifading mountant (Invitrogen). Slides were sealed with nail polish.

Samples were imaged with a Leica DM6000 DM6B fluorescence microscope (Leica) equipped with DFC9000 GT sCMOS camera using a HC PLAN APO 100x/1.4 OIL PH3 objective, using the LAS X software. The filter set for imaging DAPI is the DAPI ET filter (n. 11504203, Leica), with excitation filter 350/50 (band pass), long pass dichroic mirror 400 and emission filter 460/50 (band pass). For imaging of Oregon Green the filter L5 ET was used (n. 11504166, Leica), with excitation filter 480/40, dichroic mirror 505 and emission filter 527/30.

Data was analyzed with MicrobeJ package version 5.12d [90] with ImageJ 1.52d software [91]. Recognition of cells was limited to 2 - 16 µm length. For the nucleoid and Halotag detection the nucleoid feature was used for the nucleoid length and fluorescent analysis. Cells with more than 2 identified nucleoids and defective detection were excluded from analysis. Statistical analysis was performed with MicrobeJ package version 5.12d [90].

### In-gel fluorescence

*C. difficile* strains were cultured in BHI/YE, and when appropriate induced at an OD_600_ of 0.3-0.4 with 200 ng/mL ATc concentrations for up to 3 hours. Samples were collected and centrifuged at 4°C. Pellets were resuspended in PBS and lysed by French Press. Samples were incubated with 150 nM Oregon Green substrate for HaloTag (Promega) for 30 minutes at 37°C. Loading buffer (250 mM Tris-Cl pH 6.8, 10 % SDS, 10% β-mercaptoethanol, 50% glycerol, 0.1% bromophenol blue) was added to the samples without boiling and samples were run on 12% SDS-PAGE gels. Gels were imaged with Uvitec Alliance Q9 Advanced machine (Uvitec) with F-535 filter (460 nm).

### Spot assay

Cells were grown until OD_600_ of 1.0 in BHI/YE. The cultures were serially diluted (10^0^ to 10^-5^) and 2 µL from each dilution were spotted on BHI/YE supplemented with CDSS, thiamphenicol and 200 ng/mL ATc when appropriate. Plates were imaged after 24 hours incubation at 37ºC.

### Tethered Particle Motion measurements

A dsDNA fragment of 685bp with 32% G+C content (sso685) was used for Tethered Particle Motion experiments. This substrate was generated by PCR using the forward biotin-labelled primer Sso10a-2Nde and the reverse digoxygenin (DIG) labelled primer Sso10a-2Bam685 from pRD118 as previously described [92]. The PCR product was purified using the GenElute PCR Clean-up kit (Sigma-Aldrich).

Tethered Particle Motion (TPM) measurements were done as described previously [77, 78] with minor modifications. In short, anti-digoxygenin (20 µg/mL) was flushed into the flow cell and incubated for 10 minutes to allow the anti-digoxygenin to attach to the glass surface. To block unspecific binding to the glass surface, the flow cell was incubated with BSA and BGB (Blotting grade Blocker) in buffer A (10 mM Tris (pH 7.5), 150 mM NaCl, 1 mM EDTA, 1 mM DTT, 3% glycerol, and 100 μg/mL acetylated BSA, 0.4% BGB) for 10 minutes. To tether DNA to the surface, DNA (labeled with Biotin and DIG) diluted in buffer B (10 mM Tris (pH 7.5), 150 mM NaCl, 1 mM EDTA, 1 mM DTT, 3% glycerol, and 100 μg/mL acetylated BSA) was flushed into the flow cell and incubated for 10 minutes. Streptavidin-coated polystyrene beads (0,44 μm in diameter) diluted in buffer B were introduced into the sample chamber and incubated for at least 10 minutes to allow binding to the biotin-labelled DNA ends. Before flushing in the protein in buffer C (20 mM HEPES (pH 7.9), 60 mM KCl, and 0.2% (w/v) BGB), the flow cell was washed twice with buffer C to remove free beads. Finally the flow cell was sealed, followed by incubation with protein or experimental buffer for 10 minutes. The measurements were started after 6 minutes of further incubation of the flow cell at a constant temperature of 25 °C. More than 300 beads were measured for each individual experiment. All experiments were performed at least in duplicate.

The analysis of the TPM data was performed as previously described [78]. Equation 1 was used to calculate the RMS of the individual beads.

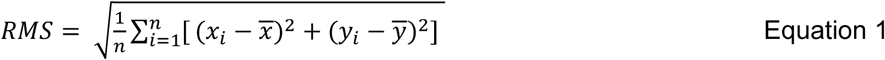

where x and y are the coordinates of the beads, 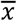 and 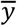 are averaged over the full-time trace. The RMS value of each measured condition was acquired by fitting a Gaussian to the histogram of the RMS values of individual beads.

All the pictures were prepared for publication in CorelDRAW X8 (Corel).

## Acknowledgements

Work in the group of WKS is supported by a Vidi Fellowship (864.10.003) of the Netherlands Organization for Scientific Research (NWO) and a Gisela Their Fellowship from the Leiden University Medical Center. Work in the group of RTD is supported by a VICI grant (016.160.613) of the Netherlands Organization for Scientific Research (NWO), Human Frontier Science Program [RGP0014/2014] and a China Scholarship Council to L.Q. [201506880001]. The authors thank Jeroen Corver for helpful discussions and Patricia Amaral for her assistance with the graphical abstract.

## Conflict of interest

WKS has performed research for Cubist. The company had no role in the design or interpretation of these experiments or the decision to publish. Others: none to declare.

## Notes

#### Summary of Updates

Provided additional controls with HupAQED mutant - including crosslinking experiments, in vivo interaction assays; added data on effects of HupA overproduction on cell viability

