## Supplemental Figures for "The bacterial chromatin protein HupA can remodel DNA and associates with the nucleoid in *Clostridium difficile*"

**Fig. S1 -** Electrostatic surface potential of *C. difficile* HupA$^{QED}$. The electrostatic potential is in eV with the range shown in the corresponding color bar.

**Fig. S2** – HupA$^{QED}$ can interact with HupA. (a) Western-blot analysis of glutaraldehyde cross-linking of HupA-HaloTag with HupA$_{6xhis}$ and HupA$^{QED}{}_{6xhis}$. 100 ng of the indicated proteins were incubated with 0%, 0.0006% and 0.006% glutaraldehyde for 30 min at room temperature. The samples were resolved by SDS-PAGE and analysed by immunoblotting with anti-his antibody. Crosslinking between HupA-HaloTag (46 kDa) and the HupA$^{(QED)}{}_{6xHis}$ monomers (11 kDa) resulted in bands corresponding to the approximate molecular weight of homodimers of HupA$^{(QED)}{}_{6xHis}$ (22 kDa), homodimers of HupA-HaloTag (92 kDa) and heterodimers (57 kDa). Additional bands of lower molecular weight HupA are observed that likely represent breakdown products and an unknown species is observed higher in the gel (both indicated with *). No difference is evident between the crosslinking with HupA$_{6xhis}$ and HupA$^{QED}{}_{6xhis}$, suggesting both can form mixed multimers with HupA-HaloTag. (b) Luciferase activity of strains AP182 ($P_{tet}$-*bitluc*$^{opt}$), AP122 ($P_{tet}$-*hupA-smbit/hupA-lgbit*), AP184 ($P_{tet}$-*smbit-hupA/lgbit*) and AP212 ($P_{tet}$-*hupA*$^{QED}$-*smbit/hupA-lgbit*). Cells were induced with 200 ng/mL anhydrotetracycline (ATc) for 60 min. Optical density-normalized luciferase activity (LU/OD) is shown right before induction (T0, blue bars) and after 1 hour of induction (T1, red bars). The averages of biological duplicate measurements are shown, with error bars indicating the standard deviation from the mean. A positive interaction was defined on the basis of the negative control (AP184) as a luciferase activity of >1000 LU/OD. No significant difference was detected at T0. At T1, only AP184 was significant different from all other samples with $p<0.0001$ (*) by two-way ANOVA.

**Fig. S3 –** Schematic representation of tethered particle motion experiments. The dsDNA molecule is labelled with digoxygenin (Dig, yellow) and biotin (orange). The dsDNA is tethered via the anti-digoxigenin antibody (anti-Dig, blue) linked to the microscope slide surface and via the streptavidin (Strep, green) linked to the bead (grey). The dotted line represents the amplitude of the bead movement. Addition of HupA leads to DNA compaction, evident as a restriction of bead movement. (b) Excursion of the bead with bare DNA (blue) and with 1600 nM HupA (red) on x-y coordinates. The root mean square (RMS) value of the excursion of each individual bead was calculated from the x- and y-coordinates, as represented by the equation.

Figure S1

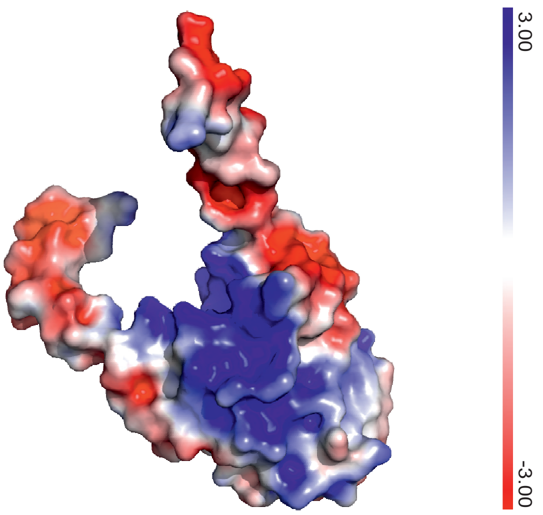

Figure S2

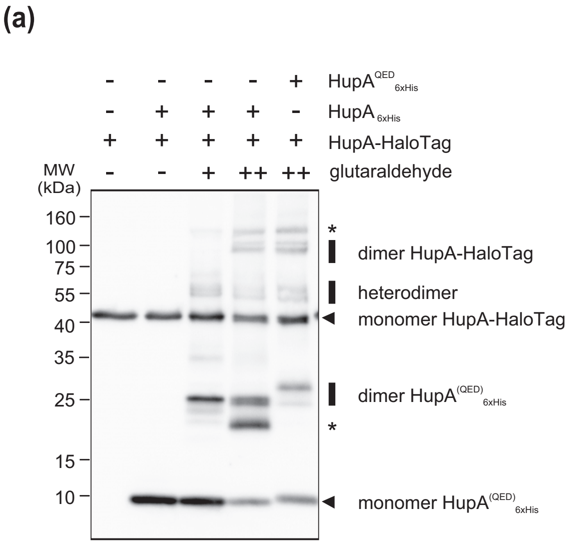

(a)
HupA^QED 6xHis
HupA 6xHis
HupA-HaloTag
glutaraldehyde
MW (kDa)
160
100
75
55
40
35
25
15
10
dimer HupA-HaloTag
heterodimer
monomer HupA-HaloTag
dimer HupA(QED) 6xHis
monomer HupA(QED) 6xHis


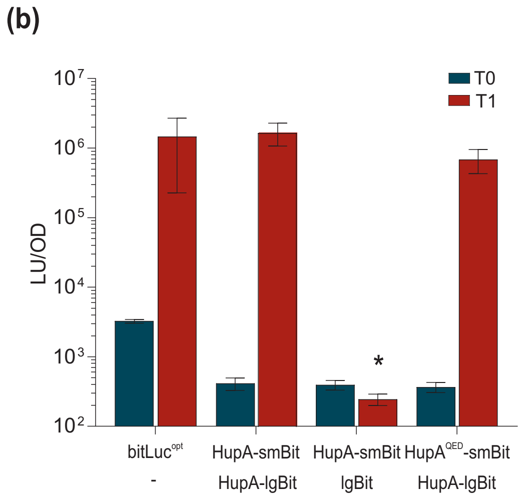

(b)
T0
T1
LU/OD
10^7
10^6
10^5
10^4
10^3
10^2
bitLuc^opt
-
HupA-smBit
HupA-lgBit
HupA-smBit
lgBit
HupA^QED-smBit
HupA-lgBit

### Figure S3

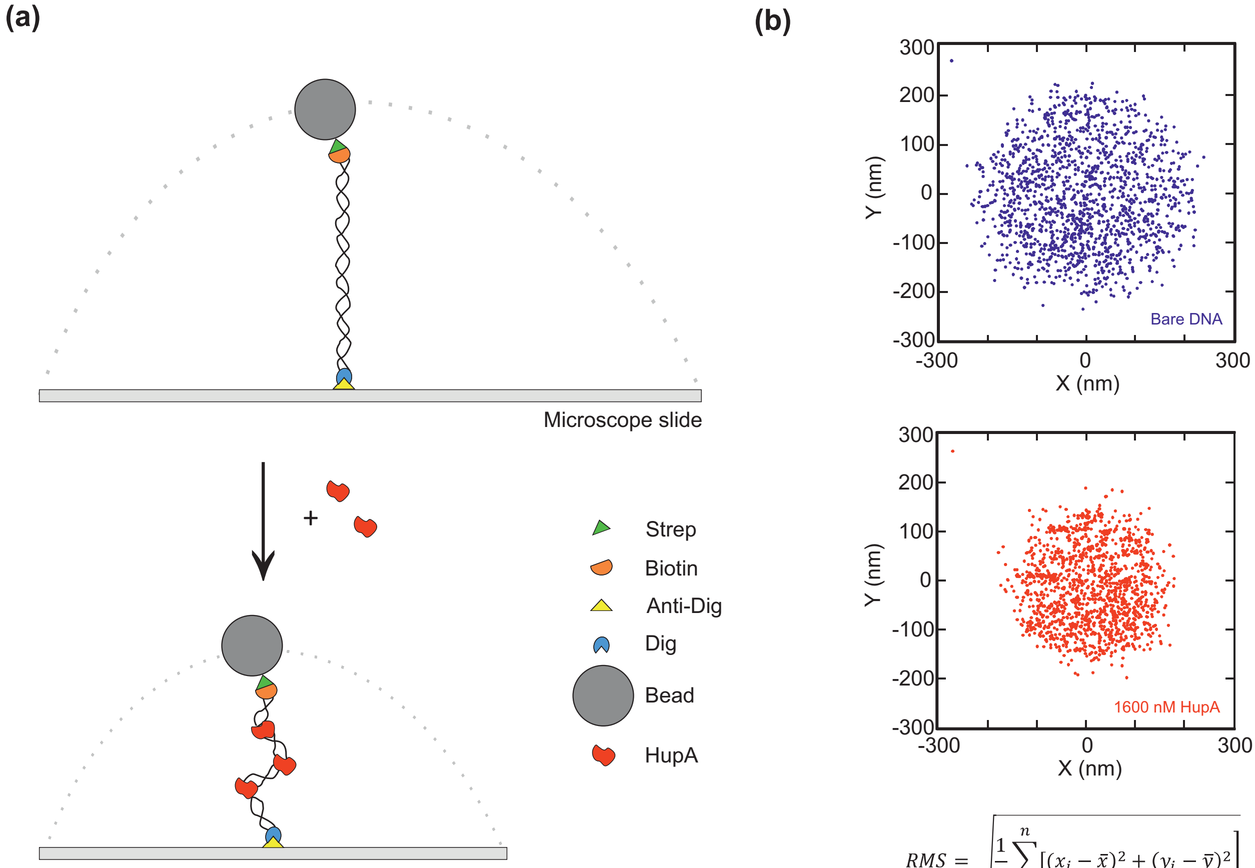


$$RMS = \sqrt{\frac{1}{n}\sum_{i=1}^{n}\left[(x_i - \bar{x})^2 + (y_i - \bar{y})^2\right]}$$
