## Supplemental Methods for "The bacterial chromatin protein HupA can remodel DNA and associates with the nucleoid in *Clostridium difficile*"

### Construction of a split-luciferase system for *C. difficile*.

We stepwise adapted our sLuc$^{opt}$ reporter [62] by 1) removing the signal sequence (resulting in an interacellular luciferase, Luc$^{opt}$), 2) introducing the mutations corresponding to the aminoacid substitutions in NanoBiT (resulting in a full lenth luciferase in which SmBiT and LgBiT are fused, bitLuc$^{opt}$) and finally, 3) the construction of a modular vector containing polycistronic construct under the control of the ATc-inducible promoter $P_{tet}$ [1].

Modules for the vectors to be constructed were synthesized dsDNA fragments based on the codon optimized sLuc$^{opt}$ sequence, but carrying the desired point mutations. Gene synthesis was performed by Integrated DNA Technologies, Inc. All primers and plasmids are listed in Tables 1 and 3. DNA sequences of the cloned DNA fragments in all recombinant plasmids were verified by sequencing.

The *hupA-smbit* and *hupA-lgbit* fragments with restriction sites to ensure modularity of the vectors were amplified with primers oAP60/oAP65 and oAP63/oAP64, respectively, cut with *Sac*I-*Bam*HI and cloned into similarly digested pRPF185, yielding plasmid pAP135 ($P_{tet}$-*hupA-smbit*) and pAP134 ($P_{tet}$-*hupA-lgbit*). To construct the plasmid harbouring an operon encoding HupA-SmBiT and HupA-LgBiT Gibson assembly was performed. The pRPF185 plasmid backbone was PCR amplified with primer set oAP58/oAP59. The *hupA-smBit* fragment was amplified from pAP135 using primers oAP60/oAP61, and the *hupA-lgBit* fragment was amplified from pAP134 using primers oAP62/oAP63. All the PCR fragments were purified and assembled at 50°C for 30 min in Gibson assembly mix [5% PEG-8000, 10 mM $MgCl_2$, 100 mM Tris-HCl pH 7.5, 10 mM DTT, 0.8 mM dNTPs, 5 mM NAD, 5.33 U/μL Taq Ligase (Qiagen), 0.005 U/μL T5 exonuclease (NEB), 0.03 U/μL Phusion polymerase (NEB)], yielding pAP118 ($P_{tet}$-*hupA-smbit*/*hupA-lgbit*). The resulting operon contains the same ribosome binding site in front of both open reading frames.

To introduce the QED mutation in pAP118, the gene encoding for *hupA*$^{QED}$ was amplified from pAF237 with primer set oAP64/oAP110. The fragment was digested with SacI/XhoI and ligated into the similarly digested pAP118, yielding pAP210.

As controls for HupA-dependency of a possible interaction, HupA-fusions were expressed from the same operon as individual luciferase domains (either SmBiT or LgBiT). To construct $P_{tet}$-*hupA-smbit/lgbit,* pAP159 was *BamH*I/*Pvu*I digested, the 511 bp fragment was gel purified, and ligated into the similarly digested pAP118 vector. As the *lgbit* lacked a characterized start codon (GTT), the first triplet was replaced with GTG (as in HupA) according to the QuikChange protocol (Stratagene) with primer set oAP98/oAP99, yielding vector pAF256.

To construct P*tet-smbit/hupA-lgBit*, pAP118 was *BamH*I/*Pvu*I digested, the 838 bp fragment was gel purified and ligated into similarly digested pAP159. As the *smbit* lacked a characterized start codon (GTT), the first triplet was replaced with GTG (as in HupA) according to the QuikChange protocol (Stratagene) with primer set oAP96/oAP97, yielding vector pAF257.

As positive controls *C. difficile* expression plasmids encoding non-secreted luciferase (Luc$^{opt}$) and a luciferase with the aminoacid substitutions (bitLuc$^{opt}$) (Promega) [61] were constructed. Luc$^{opt}$ was amplified from pAP24 [62] with primers oAP48/oAP49, digested with *BamH*I/*Sac*I and cloned into similarly digested pRPF185. As the other controls, the start codon was replaced with GTG according to the QuikChange protocol (Stratagene) with primer set oAP98/oAP99, yielding plasmid pAF254.

To construct bitLuc$^{opt}$, the *lgbit* gene was amplified from pAP134 with primers oAP54/oAP55 (Table 3) digested with *BamH*I/*Sac*I and cloned into similarly digested pRPF185. The start codon was replaced with GTG according to the QuikChange protocol (Stratagene) with primer set oAP98/oAP99, yielding plasmid pAF259.

Vectors encoding just SmBiT and LgBiT (not part of a fusion protein) were constructed as negative controls. The *lgbit* gene was amplified from pAP134 using primer oAP54/oAP63 and the fragment digested with *BamH*I/*Sac*I, and cloned into similarly digested pRPF185. The start codon was replaced with GTG according to the QuikChange protocol (Stratagene) with primer set oAP98/oAP99, yielding plasmid pAF255. The *smbit* gene was amplified from pAP135 using primer oAP66/oAP65. The fragments were *BamH*I and *Sac*I digested, and cloned into similarly digested pRPF185. As described above, the start codon was changed to GTG with primers oAP97/oAP96, yielding pAF260.

To construct the negative control plasmid harbouring both *smbit* and *lgbit* as part of the same operon, smbit and lgbit DNA fragments were generated by PCR from pAP135 using primers oAP66/oAP61 and from pAP134 using oAP67/oAP63, respectively. The fragments were fused by overlapping PCR with primers oAP66/oAP63, digested with *BamH*I/*Sac*I and cloned into similarly digested pRPF185. As described above, the start codon was changed to GTG with primers set oAP98/oAP99 and oAP97/oAP96, yielding pAF262.

It should be noted that due to the use of the GTG start codon (similar to HupA) and the fact that fusions with the luciferase domains are C-terminal, it may be necessary to adapt the start codons of the control plasmids to that of other proteins of interest when using this system.

A schematic representations of the modular vectors constructed for this study is shown in Figure SM1.

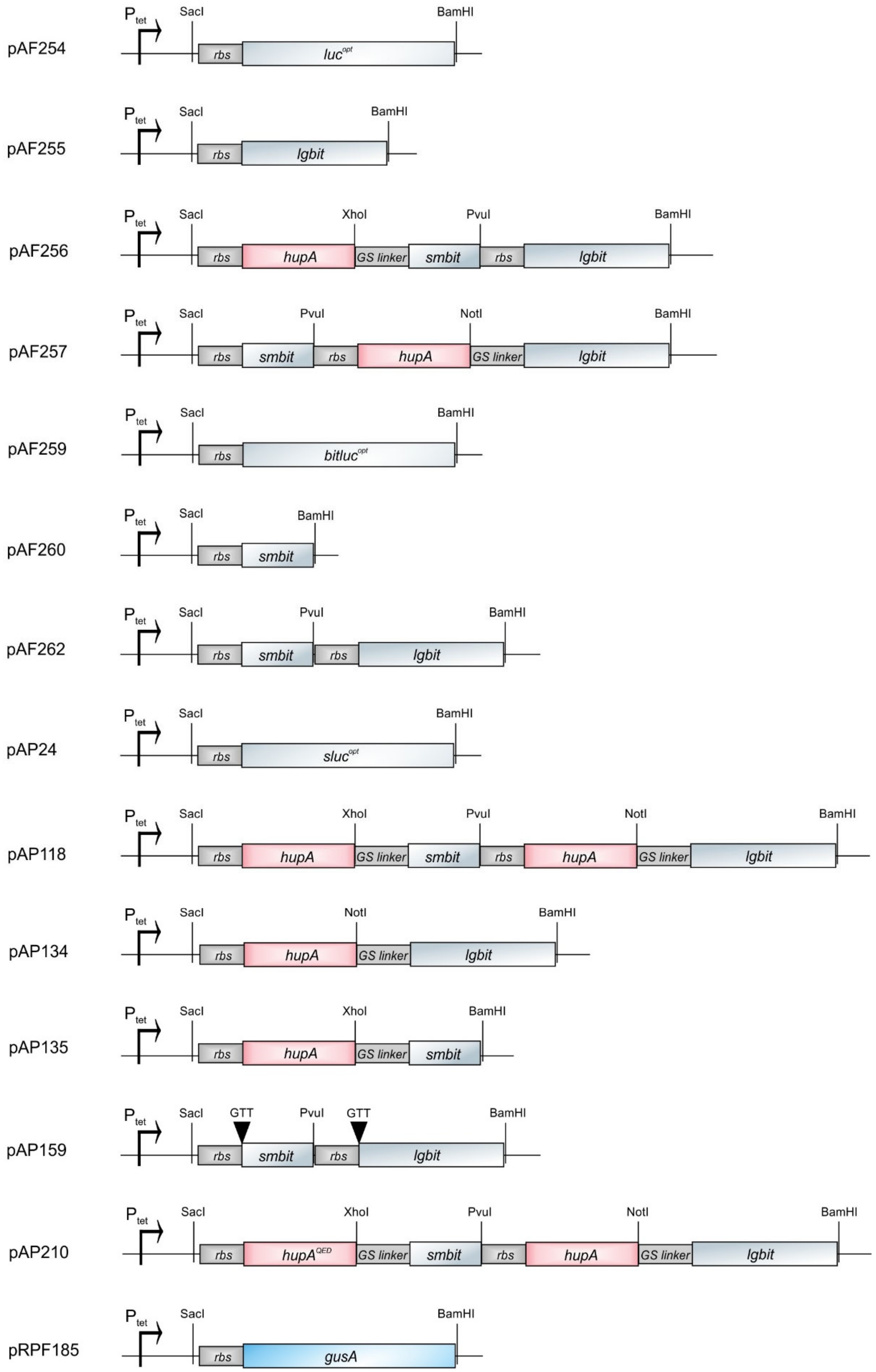


Fig. SM1 - Schematic representation of the modular vectors of the bitLuc$^{opt}$ complementation assay.

We previously described the codon optimization of the NanoLuc luciferase as part of the construction of a secreted luciferase reporter, $sLuc^{opt}$, for *C. difficile* [62]. The *C. difficile* complementation assay requires proteins to remain intracellular. Therefore, we first constructed a non-secreted luciferase reporter by removing the PPEP-1 signal sequence from $sLuc^{opt}$, yielding $Luc^{opt}$.

When expressing $sLuc^{opt}$ and $Luc^{opt}$ in *C. difficile* we detected in a significant increase of luciferase signal of the culture 1 hour after induction (Fig. SM2a). At the moment of induction $Luc^{opt}$ exhibit a luciferase signal of, slightly over the background signal of medium itself (235 ± 245 LU). After induction a significant increase in luminescence is observed for both $sLuc^{opt}$ and $Luc^{opt}$. $sLuc^{opt}$ induction results in a signal of $2.9e^{+8}$ ± $4.5e^{+7}$ LU/OD, whereas significantly lower signals were detected with $Luc^{opt}$ induction (2472785 ± 910696 LU/OD). However, the signal from an non-induced $Luc^{opt}$ (4801 ± 946 LU/OD) is lower than for the non-induced $sLuc^{opt}$ (535140 ± 44572 LU/OD), resulting in similar ~3 log increase in signal upon induction for both reporters. Thus, we conclude that the luciferase substrate can enter the cells, and the intracellular reporter is suitable for further adaptation.

For the complementation assay the substitution of specific residues was necessary to reduce spontaneous interactions between the small and large subunits of the split luciferase were necessary [2]. To determine if these substitutions potentially reduce the maximum levels of luminescence of the reconstituted luciferase, we introduced them in $Luc^{opt}$, yielding $bitLuc^{opt}$ ($Luc^{opt}$-R11E/G15A/F31L/G35A/L46R/G51A/G67A/G71A/K75E/I76V/H93P/I107L/D108N/N144T/L149M/G157S/W161Y/C164F/R166E), and assayed luminescence in the presence and absence of anhydrotetracyclin. No significant difference between $Luc^{opt}$ and $bitLuc^{opt}$ was observed (Fig. SM2b), indicating that the substitutions in $bitLuc^{opt}$ did not alter luciferase expression and activity.

Finally, we wanted to establish whether the expression of the individual subunits of the split luciferase, alone or in combination, without fusion protein would result in a positive signal in the luciferase assay. The $bitLuc^{opt}$ luciferase was split into the two subunits: lgBit

(19 kDa) and smBit (1.9 kDa). When the two subunits are expressed individually or from the same operon no significant difference was observed in the luciferase signal after induction (Fig. SM2b). This demonstrates that there is no non-specific binding of the bitLuc$^{opt}$ sub-units in *C. difficile* that could interfere with its application as a complementation assay.

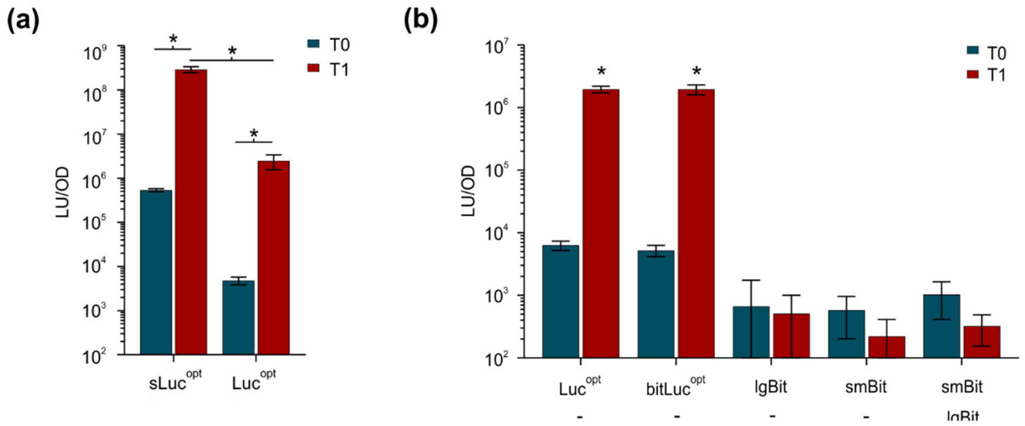


Fig. SM2 –Controls for the split luciferase (bitLuc$^{opt}$) complementation assay. Cells were induced with 200 ng/mL ATc for 60 min. Optical density-normalized luciferase activity (LU/OD) is shown right before induction (T0, blue bars) and 1 hour after induction (T1, red bars). The averages of biological triplicate measurements are shown, with error bars indicating the standard deviation from the mean. (a) Luciferase activity of AP34 ($P_{tet}$-sluc$^{opt}$; extracellular luciferase) versus AP181 ($P_{tet}$-*luc*$^{opt}$; intracellular luciferase) (b) bitLuc$^{opt}$ controls. Induction of AP181 ($P_{tet}$-*luc*$^{opt}$), AP182 ($P_{tet}$-*bitluc*$^{opt}$), AP199 ($P_{tet}$-*lgbit*), AP201 ($P_{tet}$-*smbit*) and AP202 ($P_{tet}$-*smbit/lgbit*). Interaction was defined based on the negative controls as a luciferase activity of >1000 LU/OD. Significant differences between all the values are represented * p<0,0001 by two-way ANOVA.


[1] Fagan RP, Fairweather NF. Clostridium difficile has two parallel and essential Sec secretion systems. J Biol Chem. 2011;286:27483-93.
[2] Dixon AS, Schwinn MK, Hall MP, Zimmerman K, Otto P, Lubben TH, et al. NanoLuc Complementation Reporter Optimized for Accurate Measurement of Protein Interactions in Cells. ACS Chem Biol. 2016;11:400-8.
